# Rhamnan Sulfate Reduces Atherosclerotic Plaque Formation and Vascular Inflammation

**DOI:** 10.1101/2022.02.10.479785

**Authors:** Nikita P. Patil, Almudena Gómez-Hernández, Fuming Zhang, Limary Cancel, Xu Feng, Lufeng Yan, Ke Xia, Eri Takematsu, Emily Y. Yang, Victoria Le, Megan E. Fisher, Agueda Gonzalez-Rodriguez, Carmelo Garcia-Monzon, James Tunnell, John Tarbell, Robert J. Linhardt, Aaron B. Baker

**Affiliations:** Department of Biomedical Engineering, University of Texas at Austin, Austin, TX; Department of Biochemistry and Molecular Biology, School of Pharmacy, Complutense University of Madrid, Madrid, Spain; Departments of Chemistry and Chemical Biology, Chemical and Biological Engineering, Biology, and Biomedical Engineering, Center for Biotechnology and Interdisciplinary Studies, Rensselaer Polytechnic Institute, Troy, NY; Department of Biomedical Engineering, The City College of New York, CUNY, New York, NY; Liver Research Unit, Hospital Universitario Santa Cristina, Instituto de Investigación Sanitaria Hospital Universitario de La Princesa, CIBERehd, Madrid, Spain; Institute for Cellular and Molecular Biology, University of Texas at Austin, Austin, TX; The Institute for Computational Engineering and Sciences, University of Texas at Austin, Austin, TX; Institute for Biomaterials, Drug Delivery and Regenerative Medicine, University of Texas at Austin, Austin, TX

**Keywords:** rhamnan sulfate, marine polysaccharides, atherosclerosis, NF-κB pathway, inflammation

## Abstract

**Objective:** While lipid-lowering drugs have become a mainstay of clinical therapy these treatments only slow the progression of the disease and can have side effects. Thus, new treatment options are needed to supplement the effects of lipid lowering therapy for treating atherosclerosis. We examined the use of an inexpensive and widely available marine polysaccharide rhamnan sulfate as an oral therapeutic for limiting vascular inflammation and atherosclerosis.

**Methods and Results:** We found rhamnan sulfate enhanced the barrier function of endothelial cells, preventing the deposition of LDL and maintaining barrier function even in the presence of glycocalyx-degrading enzymes. Rhamnan sulfate was also found to bind directly to FGF-2, PDGF-BB and NF-κB subunits with high affinity. In addition, rhamnan sulfate was a potent inhibitor of NF-κB pathway activation in endothelial cells by TNF-α. We treated ApoE^-/-^ mice with a high fat diet for 4 weeks and then an addition 9 weeks of high fat diet with or without rhamnan sulfate. Rhamnan sulfate reduced vascular inflammation and atherosclerosis in both sexes of ApoE^-/-^ mice but had a stronger therapeutic effect in female mice. Oral consumption of rhamnan sulfate induced a significant decrease in cholesterol plasma levels in female mice but not in male mice.

**Conclusions:** Rhamnan sulfate has beneficial effects in reducing inflammation, binding growth factors and NF-κB, enhancing endothelial barrier function and reducing atherosclerotic plaque formation in ApoE^-/-^ mice.

## Introduction

Cardiovascular disease continues to be a major cause of mortality worldwide, with roughly 17 million people dying from cardiovascular disease worldwide each year and an associated healthcare cost of over $500 billion in medical costs annually. Atherosclerosis is a chronic, progressive disease in which the accumulation of lipids, endothelial dysfunction and inflammatory processes within the vascular wall lead to plaque development.[1],[2] Atherosclerotic plaques underlie a many forms of vascular disease and can lead to organ ischemia, stroke and myocardial infarction. Pharmacotherapy with statins have become a mainstay of modern treatment of atherosclerosis but often only slow the inevitable progression of the disease and do not represent a cure for the vast majority of patients.[3, 4]

The endothelial glycocalyx is a layer of glycans that lines the interior of the artery and interacts with the flowing blood.[5] The endothelial glycocalyx provides an atheroprotective effect within the artery, in part by maintaining the endothelial barrier function that prevents lipid deposition, reduces inflammation, and lowers the risk of arteriothombosis.[6-8] During the progression of atherosclerotic disease, there is loss of the glycocalyx[9] and associated lipid deposition, and inflammation.[10, 11] Many of the major risk factors for atherosclerosis, including elevated C-reactive protein, hyperglycemia, oxidized lipids, and hyperlipidemia, lead to the depletion and degradation of the endothelial glycocalyx.[12-17] Major components of the glycocalyx include glycosaminoglycans, including heparan sulfate, chondroitin sulfate and hyaluronic acid. In the absence of these glycosaminoglycans, endothelial barrier function is compromised and immune cell adhesion is increased.[18-20]

Many marine product-derived polysaccharides have a structure roughly analogous to the polysaccharides in the glycocalyx and have oral bioavailability.[21, 22] Several of these compounds including laminarin sulfate and fucoidan have been explored as treatments for vascular disease and as anti-coagulants.[22] Sulfated polysaccharides specifically have exhibited anti-oxidant and anti-inflammatory properties making them potential therapeutics for atherosclerosis.[23] Rhamnan sulfate (RS) is a polysaccharide derived from green seaweeds including *Monostroma nitidum*. Rhamnan sulfate has a structure similar to endogenous glycosaminoglycans but is composed of a different backbone of polysaccharides which are primarily alpha-1,3-linked L-rhamnose residues that have sulfate groups. Thus, RS provides a rough approximation of the chemical structure of many of the glycans that compose the endogenous glycocalyx of the artery including that of heparan sulfate.[24] Rhamnan sulfate has been shown to decrease inflammation in vascular endothelial cells in vitro and inhibit hepatic lipogenesis in a zebrafish model.[25, 26] Previous studies from our group indicate that RS has some anti-inflammatory properties and oral bioavailability. These properties make RS a promising candidate as an oral treatment for atherosclerosis.

In this work, we examined the use of highly purified RS as a potential therapy for plaque development and vascular inflammation. Our work demonstrates that RS has an anti-inflammatory effect on endothelial cells, can bind to NF-κB and inhibit inflammatory signaling. In addition, we found that RS can directly bind to PDGF-BB and inhibit its effects on vascular smooth muscle cell (vSMC) migration and proliferation. We further tested RS as an oral therapy for atherosclerosis in male and female ApoE^-/-^ mice. We found that RS reduced atherosclerosis in both sexes, but it had stronger lipid lowering and anti-inflammatory effects on female mice in comparison to male mice. Taken together, our results support that RS has potential as an inexpensive oral treatment for atherosclerosis.

## Methods

### Cell culture

Human umbilical vein endothelial cells (HUVECs) were purchased from PromoCell. They were grown in MCDB-131 culture medium (Life Technologies) with EGM-2 growth factors (R&D Systems), 10% fetal bovine serum (FBS), L-glutamine and antibiotics. Human aortic smooth muscle cells (HAoSMCs) were grown in MCDB-131 culture medium (Life Technologies) with 10% FBS, L-glutamine and antibiotics. Cells were received at passage 2 and were not allowed to grow past passage 8. Human coronary artery endothelial cells (HCAEC) were purchased and grown in cell specific growth medium according to the manufacturer’s instructions (Cell Applications). All cells were grown at 37°C with 5% CO_2_.

### Rhamnan sulfate purification/analysis

Rhamnan sulfate was isolated from green seaweed (Monostroma nitidium sourced from Japan) powder using methods described previously.[27] The powder was homogenized in distilled water (1:30 seaweed:water) and extraction was performed at 100°C for three hours. The extract was centrifuged at 4700*g* for 10 min and the supernatant was collected. Ethanol was added to the supernatant to achieve 80% ethanol per total volume. Crude polysaccharide was precipitated from the supernatant with three volumes of anhydrous ethanol and dissolved in distilled water. The polysaccharide solution was dialyzed in a cellulose membrane (molecular weight cut-off of 3500 Da) against distilled water for three successive days. After dialysis, the RS was lyophilized and weighed. Molecular weight of a monomer of RS was 150 kDa, measured after purification, and the aggregate in 1 mg/mL solution was estimated to be 2.67×10^7^ kDa using DLS.

### Labeling of RS with FITC

Labeling and detection of RS with FITC was performed as described previously.[28] Briefly, the polysaccharide was activated by adding CNBr (8.33 mg/mL) to RS (20 mg/mL) with the mixture maintained at pH 11. Activated RS was desalted using a 20 cm Sephadex G-50 column in 0.2 M sodium borate (pH 8.0). The RS containing fractions were pooled and reacted with 2 mg fluoresceinamine. Gel filtration was used to separate RS-FITC from unreacted fluoresceinamine and the concentration of fluorescein determined by reading absorbance at 440 nm. Concentration of RS was determined using the phenolsulfuric acid method. Degree of addition was expressed as moles of fluorescein per mole of monosaccharide and molecules of fluorescein per polysaccharide molecule.

### Inhibition of uptake pathways

Endothelial and vascular smooth muscle cells were grown to confluence in MCDB-131 medium as described earlier. To prevent mitosis, mitomycin (1 mg/mL; Sigma-Aldrich) was added to the cells, with no treatment cells as control. To prevent uptake through the caveolin mediated endocytosis pathway, nystatin (1 mg/mL; Thermo Fisher Scientific) was added to the cells, using DMSO (1 mg/mL) for control. Pitstop 2 and pitstop 2 negative control (1 mg/mL; Abcam) were added to both cell types to prevent uptake through the clathrin mediated endocytosis pathway. Rottlerin (1 mg/mL; Santa Cruz Biotechnology) and DMSO control were used to test for inhibition of macropinocytosis. All groups received 1 mg/mL RS labeled with FITC. Cells were fixed after treatments for 24, 48 and 72 h, stained with DAPI and imaged using a confocal microscope.

### Proliferation and migration assays

To measure proliferation, cells were seeded on a fibronectin coated 96-well plate and grown to confluence. They were starved with 0.5% FBS medium for 24 hours, then treated with RS (100 or 1000 µg/mL) and growth factors (10 ng/mL PDGF-BB or FGF-2) for 72 hours. Proliferation was measured using the BrdU Cell Proliferation Assay Kit (Cell Signaling Technology). Migration was measured using the ORIS Cell Migration Assay Kit (Platypus Technologies). Cells were seeded on a fibronectin coated ORIS plate and grown to confluence. The cells were then placed in media with 0.5% FBS for 24 hours. The stoppers were removed, and the cells were treated with RS (100 or 1000 µg/mL) and growth factors (10 ng/mL PDGF-BB or FGF-2). Images of each well were captured every 24 hours for the duration of the experiment using a Cytation 5 Cell Imaging Multi-Mode Reader (BioTek).

### LDL permeability assay

Human coronary artery endothelial cells at passages 4-8 were plated onto fibronectin coated Transwell membranes (12 mm diameter, 0.4 µm pores, Corning) at a density of 0.5 × 10^4^ cells/cm^2^. Cells reached confluence within 2-3 days after plating, and experiments were carried out on monolayers 4-6 days post-plating. Immediately before the start of an experiment the media was changed to the experimental media consisting of phenol-red free basal media (Cell Applications) supplemented with 1% bovine serum albumin (BSA; Sigma-Aldrich). For studies with treatments, the HCAECs were incubated with growth media (control) or growth media containing RS at 25 μg/mL for 24 h, followed by a 2 h incubation with Heparinase III (HepIII; Ibex pharmaceuticals, Quebec, Canada) at 135 mU/mL or 1215 mU/mL in experimental media. After the 2 h HepIII treatment, HCAECs were rinsed twice with experimental media and the permeability of DiI-LDL was measured. Tumor necrosis factor alpha (TNF-α, 20 ng/mL; Sigma-Aldrich) and cycloheximide (chx, 3 mg/mL; Sigma-Aldrich) were used to induce elevated apoptosis and permeability.[29] The cells were grown for 5 days before incubation with TNF-α and chx (TNF-α/chx) in the presence or absence of the RS isoforms or heparin (100 µg/mL; Sigma-Aldrich). TNF-α/chx was removed after 3.5 h of incubation and the cells were allowed to recover for 20 h in the presence or absence of RS or heparin.

The experimental apparatus used for measurement of LDL was as described previously.[29, 30] The apparatus consists of eight Delrin® chambers, each connected to a laser excitation source and an emission detector. A Transwell filter containing the HCAEC monolayer was inserted and sealed within the transport chamber creating a luminal (top) and abluminal (bottom) compartment. At the beginning of each experiment DiI-LDL (5 mg/mL; Biomedical Technologies) was added to the luminal compartment. The fluorescent detection system was then used to measure the solute concentration in the abluminal compartment of each chamber as a function of time. Each transport experiment consisted of a one-hour equilibration period, followed by application of a 10 cm H_2_O pressure differential and data collection for one hour. The permeability was calculated as

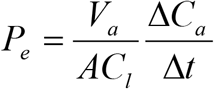

where Δ*C*_*a*_/Δ*t* is the change in the abluminal concentration with respect to time, *V*_*a*_ is the fluid volume in the abluminal compartment, *C*_*1*_ is the concentration in the luminal compartment, and *A* is the area of the filter.

### Immunostaining of heparan sulfate

Monolayers of HCAEC grown in Transwell membranes were stained for heparan sulfate using mouse monoclonal anti-heparan sulfate antibody (10E4 epitope, 1:100 in 2% goat serum, AMSBIO) following previously established protocols.[31] Briefly, monolayers were fixed in 2% paraformaldehyde and 0.1% glutaraldehyde in PBS for 30 min, blocked with 2% GS for 30 min, and incubated with primary antibody overnight at 4°C in a humidified chamber. Monolayers were then rinsed and incubated with secondary antibody Alexa Fluor 488 goat anti-mouse IgG (1:300 in 2% GS) for one hour at room temperature, and counterstained with DAPI. The stained monolayers were imaged using a ZEISS LSM 510 confocal microscope. ImageJ was used to quantify image intensity. The coverage of heparan sulfate was analyzed using the methods described in previous work.[31]

### Preparation of subendothelial matrix samples

Monolayers of HCAECs were grown for 14 days in Transwell filters before the cells were removed by a method shown to leave intact subendothelial matrix attached to the Transwell.[32] Briefly, the monolayers were washed three times in PBS, incubated in 20 mM NH_4_OH in 0.1% Triton X-100 for 5 min at room temperature, washed three times in PBS, and washed three times in basal media containing 3% BSA. The filters with subendothelial matrix were then incubated with RS for 24 h, and the permeability of the subendothelial matrix to DiI-LDL was measured.

### Nuclear fractionation and extraction of proteins

Endothelial cells were maintained in MCDB-131 medium with 0.5% FBS for at least 18h, then stimulated with TNF-α (10 ng/mL) for 30 minutes with or without RS pretreatment for 24 h (100-500 µg/mL). Tissue homogenates of endothelial cells were resuspended in a buffer, which consisted of 10 mM HEPES (pH 7.8), 15 mM KCl, 2 mM MgCl2, 0.1 mM EDTA, 1 mM dithiotheitol (DTT), and 1 mM phenylmethylsulfonyl fluoride. After 10 min on ice, the tissue homogenates were pelleted and resuspended in two volumes of the buffer. Then, 3 M KCl was added dropwise to reach a 0.39 M KCl concentration. We extracted the nuclei from the cells with incubation for 1 h at 4ºC followed by centrifuged at 12,000*g* for 1 h. The supernatants were then dialyzed in a buffer, which consisted of 50 mM HEPES (pH 7.8), 50 mM KCl, 0.1 mM EDTA, 1 mM DTT, and 1 mM phenylmethylsulfonyl fluoride with 10% (v/v) glycerol. The samples were then cleared by centrifugation and stored at −80ºC until further use. Total protein concentration was determined by BCA (Thermo Fisher Scientific). We analyzed the levels and phosphorylation of IκBα, IKKα and IKKβ in cytosolic fractions by western blotting as described below. We measured p65 in the nuclear and cytosol fractions by Western blot studies. We used β-actin and Histone H3 as control for total protein in cytosolic or nuclear fractions. The activity of NF-κB in binding DNA was assessed in nuclear fractions by a DNA Binding TransAM NF-κBp65 Assay (Active Motif).

### Western blotting

Cells were lysed using RIPA lysis buffer with added protease and phosphatase inhibitors (10 µL/mL), EDTA (10 µL/mL) and phenylmethylsulfonyl fluoride (1 mM) (Thermo Fisher Scientific). They were sonicated for three minutes and then centrifuged at 10000*g* for 10 minutes to collect total protein fraction. Concentration was determined using the Pierce BCA Protein Assay (Thermo Fisher Scientific). We performed SDS PAGE on the samples using the Invitrogen NuPAGE Bis-Tris protein gels. The proteins were transferred to nitrocellulose membranes using the wet transfer method (BioRad). The membranes were blocked with 4% StartingBlock T20 Blocking Buffer (Thermo Fisher Scientific) with 0.1% Tween-20 in PBS or TBS for one hour. The membranes were then incubated with the primary antibodies in 1% StartingBlock overnight at 4°C (**Supplemental Table 1**). The membranes were then incubated with HP conjugated secondary antibodies (Cell Signaling) at 1:3500 dilution for two hours in 1% Blotto. SuperSignal West Femto ECL solution (Thermo Fisher Scientific) was used to detect the antibodies and imaging was performed on a G:BOX imaging system (Syngene, Inc.).

### Murine model of atherosclerosis

All animal studies were performed with the approval of the University of Texas at Austin Institutional Animal Care and Use Committee (IACUC) and in accordance with NIH guidelines “Guide for Care and Use of Laboratory Animals” for animal care. We used male and female ApoE^-/-^ mice (B6.129P2-Apoe^tm1Unc^/J) for this study (Jackson Laboratories, Inc.). The animals were given high fat chow which was the standard formulation Clinton/Cybulsky high fat rodent diet with regular casein and 1.25% added cholesterol (D12108C; Research Diets, Inc.). For RS treated animals, the high fat diet (HFD) was formulated with 0.75 g of RS per kg diet. At 12 weeks of age, the mice were switched from normal chow to high fat chow for 4 weeks. The HFD group remained on this diet for an additional 9 weeks while the treatment group was switched to high fat chow with RS. At 4, and 12 weeks after start of HFD, the vessels of the mice were imaged using high resolution ultrasound as described below. At 13 weeks of HFD, the mice were sacrificed, and the blood, heart, aorta, liver and white adipose tissues were harvested for further analysis. For control, we used male and female wild type mice (n=5) fed a standard diet for 13 weeks.

### High resolution ultrasound imaging

High resolution ultrasound was performed using the Visual Sonics VEVO 2100 system with the MS500D transducer in the ApoE^-/-^ model. The mice were anesthetized with isoflurane and body temperature monitored using a probe. Prior to imaging, the fur on the chest was removed using Nair. First, the longitudinal section of the aortic arch was located using the B-mode option. Pulsed wave Doppler and 3D echocardiography images were then obtained for calculating peak systolic velocity (PSV), end diastolic velocity (EDV) and, mean velocity (MV). The longitudinal section of the left common carotid artery was located using B-mode and pulsed wave Doppler images were obtained for the same parameters in the carotid artery. Finally, the cross section of the aortic arch was imaged in B-mode and color Doppler. Diameter measurements of the aortic arch determined from cross sections were used to calculate circumferential strain using the following formula:

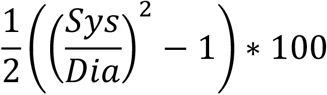

Where sys and dia are the peak systolic and end diastolic diameters.

### Histological staining

Aortic roots at optimal cutting temperature were embedded in NEG-50 and cut into 7 μm thick sections using a cryostat (CM1510 S; Leica). These sections were stained with Oil Red O/hematoxylin to measure lipid depot using the protocol below. Individual lesion area was determined by averaging the maximal values. For the liver samples, the tissues were placed in NEG-50 embedding medium and then frozen in liquid nitrogen cooled isopentane. The samples were serially sectioned to create 7 μm thick cryosections and Oil Red-O staining was performed. For the fat tissues, white adipose tissues (gonadal and inguinal depots) were fixed overnight in 4% formaldehyde and paraffin sections were created using standard methods. The size of adipocytes in white adipose tissues were quantified in H&E staining using Image J software.

### Oil Red-O staining

A stock Oil Red O (Electron Microscopy Sciences) solution was made with 3 mg/mL Oil Red O in 99% isopropanol. The stock was diluted in a 3:2 ratio in ultrapure water. Frozen cryosections were air dried and then fixed in 10% formalin. They were stained with Oil Red O and counterstained with Mayer’s hematoxylin (Electron Microscopy Sciences). The sections were mounted with aqueous mounting medium for imaging (Vector Labs).

### Immunostaining for tissues

Lesional macrophages (F4/80), vSMCs (α-SMA), PECAM-1, eNOS, IκBα and p-p65 were detected by immunoperoxidase or immunofluorescence (**Supplemental Table 1**). Positive staining was expressed as percentage of total plaque area or number of positive cells per lesion area. In each experiment, negative controls without the primary antibody or using a nonrelated antibody were included to check for nonspecific staining.

### En face imaging of aorta

Atherosclerotic lesions were quantified by en face analysis of the whole aorta. For en face preparations, the aorta was opened longitudinally, from the heart to the iliac arteries, while still attached to the heart and major branching arteries in the body. The aorta from the heart to the iliac bifurcation was then removed and was pinned out on a white wax surface in a dissecting pan using stainless steel pins 0.2 mm in diameter. After overnight fixation with 4% paraformaldehyde and a rinse in PBS, the aortas were immersed for 6 min in a filtered solution containing 0.5% Oil Red-O, 35% ethanol and 50% acetone and destained in 80% ethanol. The Oil Red-O stained aortas were photographed, and the atherosclerotic lesions were quantified using IP Win32 v4.5 software.

### Pharmacokinetics of rhamnan sulfate in vivo

To test for oral bioavailability of RS, male mice (C57BL/6; Jackson Labs) were given RS-FITC (0.25 g RS/kg mouse) though oral gavage. Mice were sacrificed at the time points of 0, 1, 4, 12 and 24 hours (n=2), and blood, aorta, liver and heart were harvested. The tissue samples were lysed and RS-FITC concentration measured by reading fluorescence intensity with correction for background fluorescence. A calibration curve was made for RS-FITC in each tissue to convert fluorescence to concentration.

To measure the half-life of RS in the blood a mouse was injected intravenously via the tail vein with 0.5 mg of FITC conjugated RS (FITC-RS). A 10 ml blood sample was collected from the saphenous vein 1 minute after injection, and then every five minutes for 25 minutes. The blood samples were centrifuged at 12,000 rpm for 5 minutes and the plasma was used for fluorescence measurements. Fluorescence intensity was measured using a plate reader (BioTek). A calibration curve for FITC-RS in mouse plasma was created which allowed the conversion of fluorescence intensity into concentration. The plasma concentration data was fitted to an exponential (first order elimination process is assumed). Extrapolating from the resulting equation we estimate the concentration at t=0 is 0.278 mg/mL. The apparent volume of distribution, V_D_, is calculated from the relationship V_D_ = Dose/C_p_^0^, where C_p_^0^ is the concentration at t=0. We calculated V_D_ to be 1.8mL. This is approximately equal to the blood volume (the blood volume of a mouse is 77-80 mL/g). The half-life was calculated from the elimination rate constant 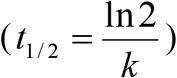. With the data above, (k=0.0328 min^-1^) the half-life is 21.1 minutes. Extrapolating from the data, the concentration drops below the lowest effective concentration we have tested (1 mg/mL) after 170 minutes.

### Raman spectroscopy

Raman imaging was performed using a custom built 830nm confocal Raman microscope with a 60x water immersion objective (NA = 1.2; Olympus).[33] The scattered light was collected by a spectrograph and a CCD camera though a 50 μm core diameter fiber, which also acts as the pinhole of the confocal system. Fresh sections with 10 μm thickness were mounted on low-background Raman substrates (magnesium fluoride window or quartz slide) for Raman imaging. For every animal, one liver section was prepared, and 2 – 5 regions of interest were randomly selected from each section. Raman imaging was performed with a 0.25 or 0.5 s integration time and a step size of 0.75 or 1 μm. Image size varied from 24 × 24 μm^2^ to 40 × 40 μm^2^. Data preprocessing was performed using MATLAB (R2017a, MathWorks). Preprocessing steps included wavenumber calibration, background removal, cosmic ray removal, smoothing, and fluorescence background removal. Spectra were normalized to the area under the curve between 600 and 1800 cm^-1^. Cluster analysis was performed for each image by k-means algorithm.[34] The first 100 principal components accounting for 95% - 99% of the variation in the data set served as the input for the k-means. Each image contained at most three clusters, annotated as the lipid-rich zone, protein-rich zone, and others. Only the spectra within the lipid-rich zone were extracted to calculate the degree of unsaturation. The average Raman spectrum of the lipid-rich zone was compared between different groups. The degree of unsaturation was calculated by the peak ratio of 1656 cm^-1^ (integrated from 1645 – 1675 cm^-1^) to 1441 cm^-1^ (integrated from 1420 – 1480 cm^-1^), which corresponds to the ratio of C=C stretching vibration and the band related to the CH_2_ scissoring mode.[34, 35]

### Preparation of rhamnan sulfate and heparin biochip

Biotinylated RS or heparin was prepared by conjugating the reducing end to amine-PEG3-Biotin (Pierce).[36] In brief, RS or heparin (2 mg) and amine-PEG3-Biotin (2 mg, Pierce) were dissolved in 200 µl H_2_O and 10 mg NaCNBH_3_ was added. The reaction mixture was heated at 70 °C for 24 h, after that a further 10 mg NaCNBH_3_ was added and the reaction was heated at 70 °C for another 24 h. After cooling to room temperature, the mixture was desalted with the spin column (3,000 MWCO). Biotinylated RS or heparin was collected, freeze-dried and used for surface plasmon resonance (SPR) chip preparation. The biotinylated RS or heparin was immobilized to a streptavidin chip (GE Healthcare) based on the manufacturer’s protocol. Successful immobilization of RS or heparin was confirmed by the observation of about 100 resonance unit (RU) increase on the sensor surface after 20 µl injection of RS or heparin (1 mg/mL). The control flow cell was prepared by 1 min injection with saturated biotin.

### Surface plasmon resonance of binding between rhamnan sulfate and proteins

Recombinant human FGF-1 and FGF-2 were a gift from Amgen. We purchased human antithrombin III (AT) (Hyphen Biomed), recombinant human platelet-derived growth factor (PDGF-BB), NF-κB p50, and NF-κB p65 (Abcam). The interactions between RS and proteins were measured using the BIAcore 3000 SPR system (GE Healthcare). The protein samples were diluted in HBS-EP buffer (0.01 M HEPES, 0.15 M NaCl, 3 mM EDTA, 0.005% surfactant P20, pH 7.4). Different dilutions of protein samples were injected at a flow rate of 30 µL/min. At the end of the sample injection, HBS-EP buffer was flowed over the sensor surface to facilitate dissociation. After a 3 min dissociation time, 30 µL of 2 M NaCl was injected to fully regenerate the surface. The response was monitored as a function of time (sensorgram) at 25°C.

### Statistical analysis

All results are shown as mean ± standard error of the mean. Comparisons between only two groups were performed using a 2-tailed Student’s t-test. Multiple comparisons between groups were analyzed by 2-way ANOVA followed by a Tukey post-hoc test. A 2-tailed probability value *p* < 0.05 was considered statistically significant.

## Results

### Rhamnan sulfate is internalized by macropinocytosis and is dependent on proliferation on vSMCs

To examine the kinetics of uptake of RS by vascular cells, we incubated endothelial cells and aortic vSMCs with FITC-labeled RS. We found that RS was detectable after 24 hours and that there was nuclear fluorescence from the RS after 48 hours (**Fig. 1A, B**). Based on the timing of the nuclear localization of RS, we hypothesized that RS was entering the nucleus during the disassembly and reassembly of the nuclear envelope during cell division. To test this hypothesis, we mitotically arrested cells using mitomycin and measures RS uptake. In vSMCs, mitomycin significantly inhibited the uptake and nuclear localization of RS (**Fig. 1C**). However, this effect was not seen in endothelial cells (**Fig. 1D**). We also treated the cells with inhibitors of caveolin-mediated endocytosis (nystatin) and clathrin-mediated endocytosis (Pitstop 2) but neither had clear inhibitory effects on RS uptake (**Fig. 1E-H**). In contrast, treatment with an inhibitor of macropinocytosis (rottlerin) significantly inhibited uptake of RS in both endothelial cells and vSMCs (**Fig. 1I, J**).

**Figure 1.**
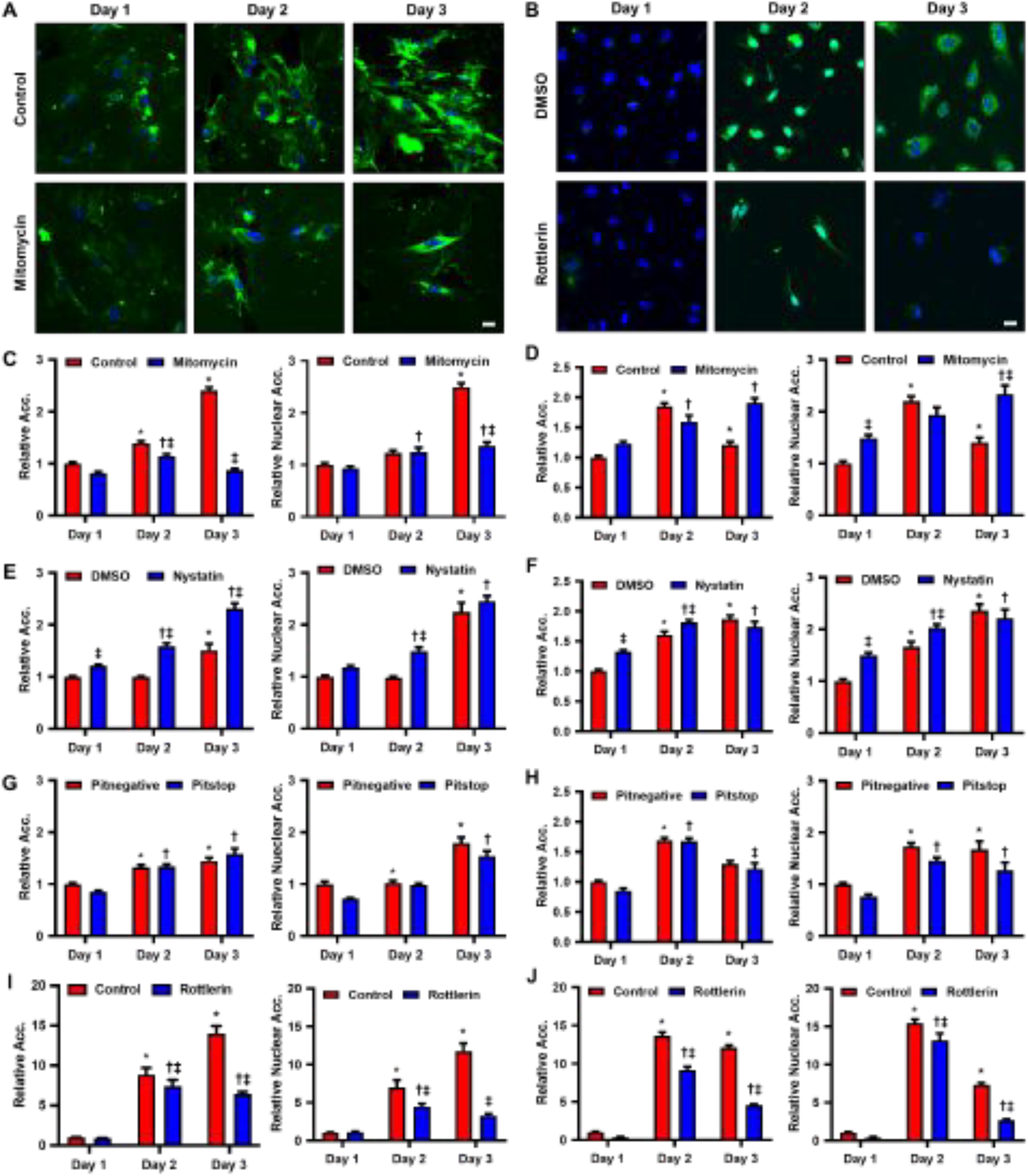
Vascular cells internalize RS primarily through macropinocytosis. (A) Vascular smooth muscle cells were incubated with FITC-labeled RS (1000 μg/mL) with or without treatment with mitomycin for the indicated times. Scale bar = 20 μm. (B) Endothelial cells were incubated with FITC-labeled RS (1000 μg/mL) with or without treatment with rottlerin. Scale bar = 20 μm. C) Uptake of RS in (s treated with mitomycin decreased over three days. \**p <* 0.05 versus control day 1. ^†^*p <* 0.05 versus treatment group on day 1. ^‡^*p <* 0.05 versus control for each time point. D) In endothelial cells treated with mitomycin, uptake of RS appeared to increase over three days. \**p <* 0.05 versus control day 1. ^†^*p <* 0.05 versus treatment group on day 1. ^‡^*p <* 0.05 versus control for each time point. Quantification of total and nuclear uptake of RS in (E) vSMCs or (F) endothelial cells treated with nystatin, an inhibitor of caveolin-mediated endocytosis. Quantification of total and nuclear uptake of RS in (G) vSMCs or (H) endothelial cells treated with pitstop 2, an inhibitor of clathrin-mediated endocytosis. Quantification of total and nuclear uptake of RS in (I) vSMCs or (J) endothelial cells treated with rottlerin, an inhibitor of macropinocytosis. \**p <* 0.05 versus control day 1. ^†^*p <* 0.05 versus treatment group on day 1. ^‡^*p <* 0.05 versus control for each time point. (F) Images of endothelial cells that were incubated with FITC-labeled RS (1000 μg/mL) with or without treatment with rottlerin for the indicated times. Scale bar = 20 μm. Quantification of total and nuclear uptake of RS in endothelial cells treated with (B) mitomycin, (C) nystatin, (D) pitstop, and (E) rottlerin. \**p <* 0.05 versus control day 1. ^†^*p <* 0.05 versus treatment group on day 1. ^‡^*p <* 0.05 versus control for each time point (n = 10 for all groups).

### Rhamnan sulfate decreases proliferation and migration of vascular cells

The proliferation and migration of vSMCs in atherosclerosis and vascular injury is an important mechanism in the development of intimal hyperplasia and remodeling of atherosclerotic plaques.[37] Endothelial proliferation and migration can be beneficial in vascular healing and reendothelialization following endothelial denudation.[38] However, these processes are also important for plaque neovascularization, which may have a role in thromboembolism and plaque destabilization.[39] We treated endothelial cell and vSMCs with RS and assayed their proliferation and migration in response to growth factors. Treatment with RS significantly reduced proliferation and migration of endothelial cells treated with FGF-2 (**Fig. 2A-C**). In vSMCs, RS decreased proliferation and migration in response to PDGF-BB (**Fig. 2D-F**). We also found that RS binds with high affinity to FGF-2 (K_D_=2.6×10^−8^ M) and PDGF-BB (K_D_=2.8×10^−8^ M; **Supplemental Fig. 1**; **Supplemental Table 2**). Proliferation and migration decreased in vSMCs treated with RS and FGF-2 (**Fig. 2G, H**).

**Figure 2.**
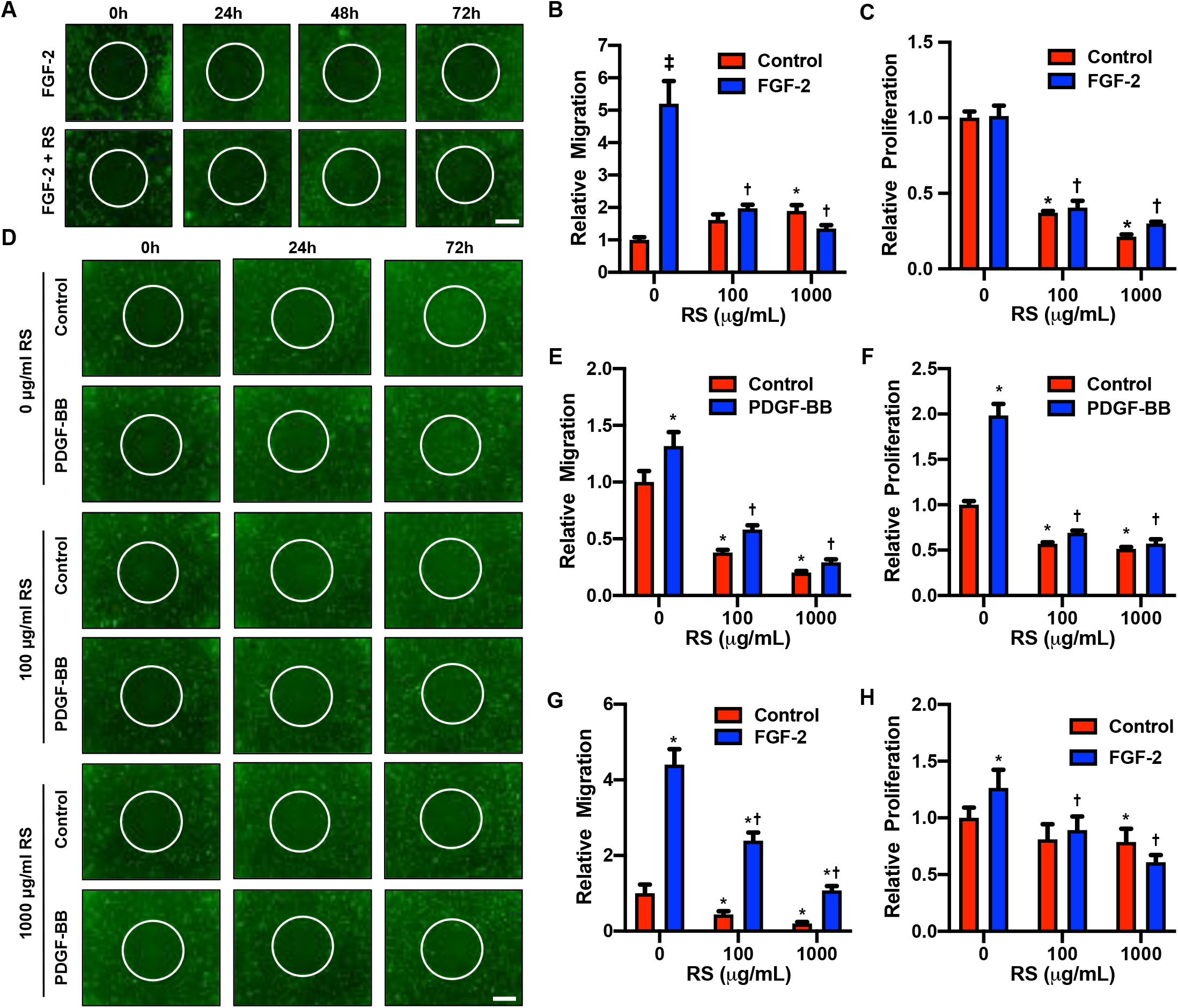
Rhamnan sulfate decreases proliferation and migration of vascular cells. (A) Endothelial cells were treated with FGF-2 and RS in ORIS assay. Scale bar = 100 μm. (B) Migration of endothelial cells treated with FGF-2 was reduced by RS (n = 10). \**p <* 0.05 versus 0 μg/mL RS, ^†^*p <* 0.05 versus 0 μg/mL RS+FGF-2, ^‡^*p <* 0.05 versus respective control. (C) Proliferation of endothelial cells treated with FGF-2 and RS decreased after 72 hours (n = 16). \**p <* 0.05 versus 0 μg/mL RS, ^†^*p <* 0.05 versus 0 μg/mL RS+FGF-2, ^‡^*p <* 0.05 versus respective control. (D) Vascular smooth muscle cells expressing GFP were treated with PDGF-BB and 0, 100 and 1000 μg/mL of RS in ORIS assay. Scale bar = 100 μm. (E) Migration of vSMCs treated with PDGF-BB was decreased by RS (n = 10). \**p <* 0.05 versus 0 μg/mL RS, ^†^*p <* 0.05 versus 0 μg/mL RS+PDGF-BB. (F) Vascular smooth muscle cells treated with PDGF-BB and RS proliferated less after 72 h (n = 16). \**p <* 0.05 versus 0 μg/mL RS, ^†^*p <* 0.05 versus 0 μg/mL RS+PDGF-BB, ^‡^*p <* 0.05 versus respective control. (G) Migration of cells vSMCs treated with FGF-2 and RS increased after 72 hours of treatment (n = 10). \**p <* 0.05 versus 0 μg/mL RS, ^†^*p <* 0.05 versus 0 μg/mL RS+FGF-2, ^‡^*p <* 0.05 versus respective control. (H) Proliferation of vSMCs induced by FGF-2, was also reduced after 72 h with RS treatment (n = 16). \**p <* 0.05 versus 0 μg/mL RS, ^†^*p <* 0.05 versus 0 μg/mL RS+FGF-2, ^‡^*p <* 0.05 versus respective control.

### Rhamnan sulfate enhances endothelial barrier function

We next examined whether RS could alter endothelial barrier function to LDL in the presence of treatments that induce inflammation or simulate the destruction of the glycocalyx. We treated endothelial cells with heparinase III, a bacterial enzyme that degrades heparan sulfate. In untreated endothelial cells this led to a drop in heparan sulfate coverage of the cells from 96% to 36% of the cells (**Fig. 3A, B**). With RS treatment, this heparinase-induced reduction in coverage was reduced to 59%, indicating that RS prevented the degradation of endogenous heparan sulfate glycosaminoglycans (**Fig. 3A, B**). The permeability of the endothelium to LDL was lower in endothelial cells treated with RS, regardless of dose of heparinase applied (**Fig. 3C**). Using Transwell filters as the base, we also tested the effect of 30 minutes of RS incubation on the blank filter, subendothelial matrix and endothelial monolayer. Treatment with RS reduced LDL permeability by 2.7-fold for the blank filter, 8-fold for the subendothelial matrix and 4.4-fold for the endothelial monolayer (**Fig. 3D**). These results indicated that RS accumulates both in the matrix and on the endothelial monolayer, where it can provide enhanced barrier function. We also tested the effectiveness of RS in reducing LDL permeability in endothelial cells after combined treatment with TNF-α and the protein synthesis inhibitor cycloheximide (CHX). While LDL permeability increased after the TNF-α/CHX treatment in control cells, it remained at basal levels with RS incubation (**Fig. 3E**). In contrast we did not observe this effect with treatment with heparin (**Fig. 3E**). We also examined the permeability of endothelial monolayers to VLDL and found that it was lower with RS treatment (**Fig. 3F**). These results suggest that RS can reduce endothelial permeability to LDL and improve overall barrier function.

**Figure 3.**
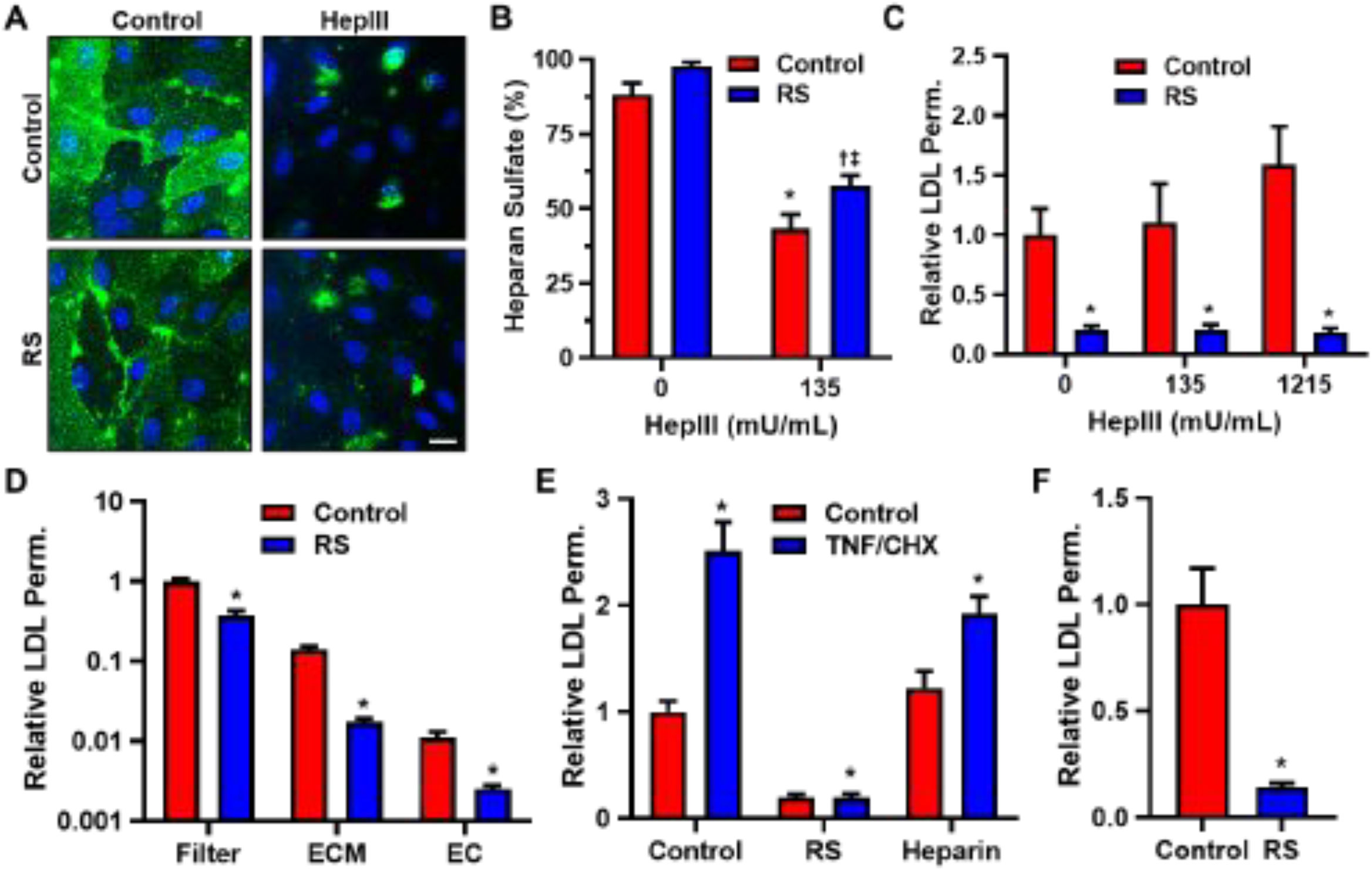
Rhamnan sulfate reduces LDL permeability in endothelial cells. (A) Endothelial cells were treated with RS (25 μg/mL) and heparinase III (135 mU/mL) and immunostained for heparan sulfate. Scale bar = 20 μm. (B) Heparan sulfate coverage, reduced by heparinase III, was increased in ECs treated with RS (n = 8). \**p <* 0.05 versus 0 mU/mL heparinase III, ^†^*p <* 0.05 versus 0 mU/mL heparinase III+RS, ^‡^*p <* 0.05 versus 135 mU/mL heparinase III. (C) Permeability of LDL was decreased in ECs treated with RS regardless of heparinase III dose (n = 4-8). \**p <* 0.05 versus respective control. (D) Permeability of LDL was reduced in blank Transwell filters and those with subendothelial matrix and endothelial cell monolayer with RS incubation (n = 4). \**p <* 0.05 versus respective control. (E) Permeability of LDL was lower with RS treatment despite addition of TNF-α and cycloheximide. Heparin treatment produced no change in the same conditions (n = 4-19). \**p <* 0.05 versus control, ^†^*p <* 0.05 versus control+TNF-α/chx. (F) Permeability of VLDL also decreased by RS treatment (n = 8). \**p <* 0.05 versus control.

### Rhamnan sulfate reduces TNF-α induced NF-κB activation in human endothelial cells

The NF-κB pathway is an important pathway in controlling endothelial inflammation and is implicated in atherosclerosis, lipid metabolism and control of vSMC proliferation.[40, 41] In the canonical NF-κB pathway, an NF-κB dimer is sequestered in the cytoplasm though interaction with a member of the IκB family of proteins (eg. IκBα). Activation of a receptor leads to the recruitment of adaptor proteins followed by recruitment of the IκB kinase complex (IKK; typically consisting of IKKα, IKKβ and IKKγ), which subsequently phosphorylates IκB and eventually leads to its degradation. The destruction of IκB allows NF-κB dimers to translocate to the nucleus where they can control gene transcription. To examine if RS could alter signaling though the NF-κB pathway, we treated endothelial cells with TNF-α for varying times and measured the activation of NF-κB pathway signaling intermediates using western blotting. We found that TNF-α induced an increase in IκBα phosphorylation after 10 min of treatment, indicating activation of the NF-κB pathway and that treatment with RS reduced this response (**Fig. 4A, B**). After 30 minutes of TNF-α treatment there was increased IκBα degradation and this was not affected by RS but basal levels of IκBα were increased (**Fig. 4C, D**). We also found that NF-κBp65 protein levels significantly decreased by RS treatment in the nuclear fraction and increased in the cytoplasm fraction after 30 minutes of TNF-α treatment, indicating reduced activation of the NF-κB pathway (**Fig. 4E, F; Supplemental Fig. 2**). In addition, phosphorylation of IKK, the complex responsible for the phosphorylation of IκBα, was also significantly reduced by RS treatment (**Fig. 4G, H**). Together, these findings indicate that RS affects multiple steps in the canonical pathway of NF-κB activation by TNF-α. To further confirm these findings, we measured NF-κBp65 activity directly using a TransAM assay and found that activity was reduced in the nuclear extracts of TNF-α treated endothelial cells (**Fig. 4I**). As we had observed nuclear localization of RS in our trafficking studies, we hypothesized that RS may bind directly to some of the components of the NF-κB pathway. Using surface plasmon resonance, we found that RS binds with high affinity to NF-κBp50 (K_D_= 1.1×10^−7^ M) and NF-κBp65 (K_D_= 2.3×10^−8^ M; **Fig. 4J, K**). The dissociation constant for the proteins for binding to RS was comparable to binding with heparin (**Supplemental Table 2**).

**Figure 4.**
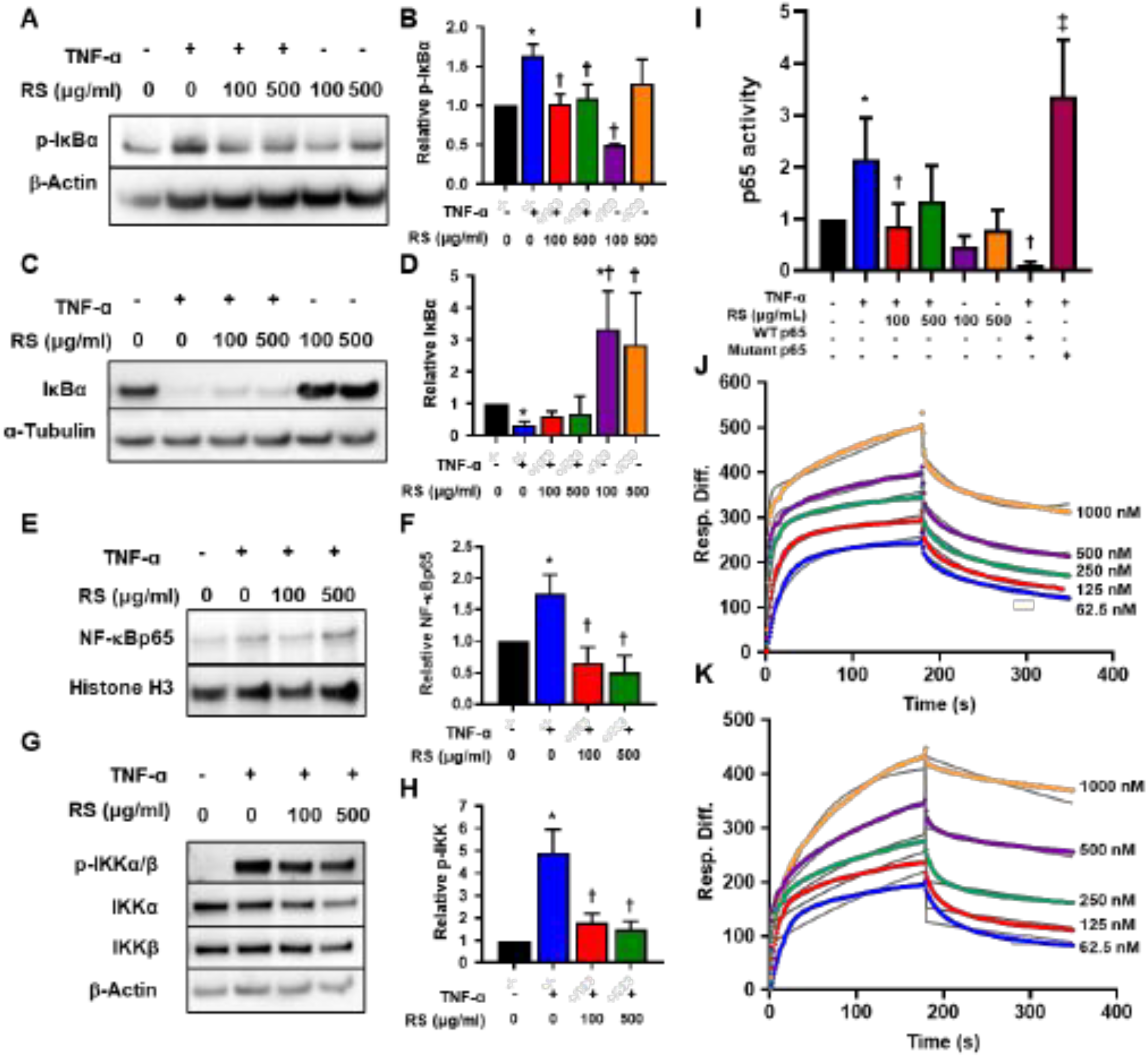
Rhamnan sulfate reduces NF-κB activation in endothelial cells. (A) Endothelial cells were treated with TNF-α (10 ng/mL) for 10 minutes to cause IκBα phosphorylation. (B) Western blot analysis showed decrease in p-IκBα in total protein of cells pre-treated with RS (n = 2-15). \**p <* 0.05 versus control, ^†^*p <* 0.05 versus TNF-α. (C) Endothelial cells were treated with TNF-α (10 ng/mL) for 30 minutes to cause degradation of IκBα phosphorylation. (D) Western blotting showed increase in IκBα in the absence of TNF-α with RS treatment (n = 7-11). \**p <* 0.05 versus control, ^†^*p <* 0.05 versus TNF-α. (E) Endothelial cells were treated with TNF-α (10 ng/mL) for 30 minutes to cause NF-κB translocation into the nucleus. (F) Western blotting showed decrease in p65 subunit of NF-κB in nuclear fraction of cells pre-treated with RS (n = 6). \**p <* 0.05 versus control, ^†^*p <* 0.05 versus TNF-α. (G) Endothelial cells were treated with TNF-α (10 ng/mL) for 10 minutes to cause activation of the IKK complexes. (H) Western blotting showed decrease in p-IKKα/β but no change in IKKα and IKKβ in total protein of cells pre-treated with RS (n = 2-11). \**p <* 0.05 versus control, ^†^*p <* 0.05 versus TNF-α. (I) TransAM assay showed reduced NF-κB/p65 activity after treatment with 100 µg/mL RS dose (n = 8). \**p <* 0.05 versus control, ^†^*p <* 0.05 versus TNF-α, ^‡^*p <* 0.05 versus WT p65. Dilutions of (J) NF-κB/p65 and (K) NF-κB/p50 were introduced on a sensor surface with RS and dissociation with HBS-EP buffer measured for three minutes. After three minutes, the surface was regenerated with 2 M NaCl and total response monitored in sensorgram (n = 5).

### Rhamnan sulfate has oral bioavailability and is found in vascular tissues after oral administration

Prior to conducting an *in vivo* experiment with oral RS, we studied the pharmacokinetics of the drug to calculate the clearance rate and uptake by aorta, heart, blood and liver. We introduced RS in mice though oral gavage and measured the concentration in tissues over 24 hours. In the abdominal aorta, RS concentration started decreasing after 4 hours but continued to rise steadily for 24 hours in the thoracic aorta (**Supplemental Fig. 3A**). In the heart and total blood plasma, RS concentrations increased up to 4 and 12 hours respectively before being cleared (**Supplemental Fig. 3B**). In the liver, there was an initial influx of RS at 4 hours and then a gradual increase in concentration after 12 hours (**Supplemental Fig. 3B**). We also injected FITC labeled RS into the blood directly and examined its clearance. We calculated the half-life of RS to be approximately 21.1 minutes in the blood following intravenous injection (**Supplemental Fig. 3C**).

### Orally administered rhamnan sulfate decreases plaque deposition in ApoE^-/-^ mice on a high fat diet

To test the effect of RS on the progression of atherosclerosis, ApoE^-/-^ mice were fed a high fat diet or a high fat diet supplemented with RS for 13 weeks (**Supplemental Fig. 4**). Blood cholesterol levels were decreased significantly in female ApoE^-/-^ mice treated with RS but not in male mice (**Fig. 4A**). There was no change in plasma triglyceride levels in both male and female mice (**Supplemental Fig. 5**). In en face preparations of the aorta, lipid deposition was decreased by 45.2% in female and 36.4% in male mice whole aortas with RS treatment in comparison to the ApoE^-/-^ HFD group (**Fig. 5B, C**). In the aortic arch, there was also significant decrease in lipid deposition in the male and female ApoE^-/-^ mice with RS treatment. Stenosis decreased in female mice only (**Fig. 5D, E**). In the thoracic aorta, there was a reduction in lipid deposition, lesion area and stenosis for female mice treated with RS but not for male mice (**Fig. 5F, G**).

**Figure 5.**
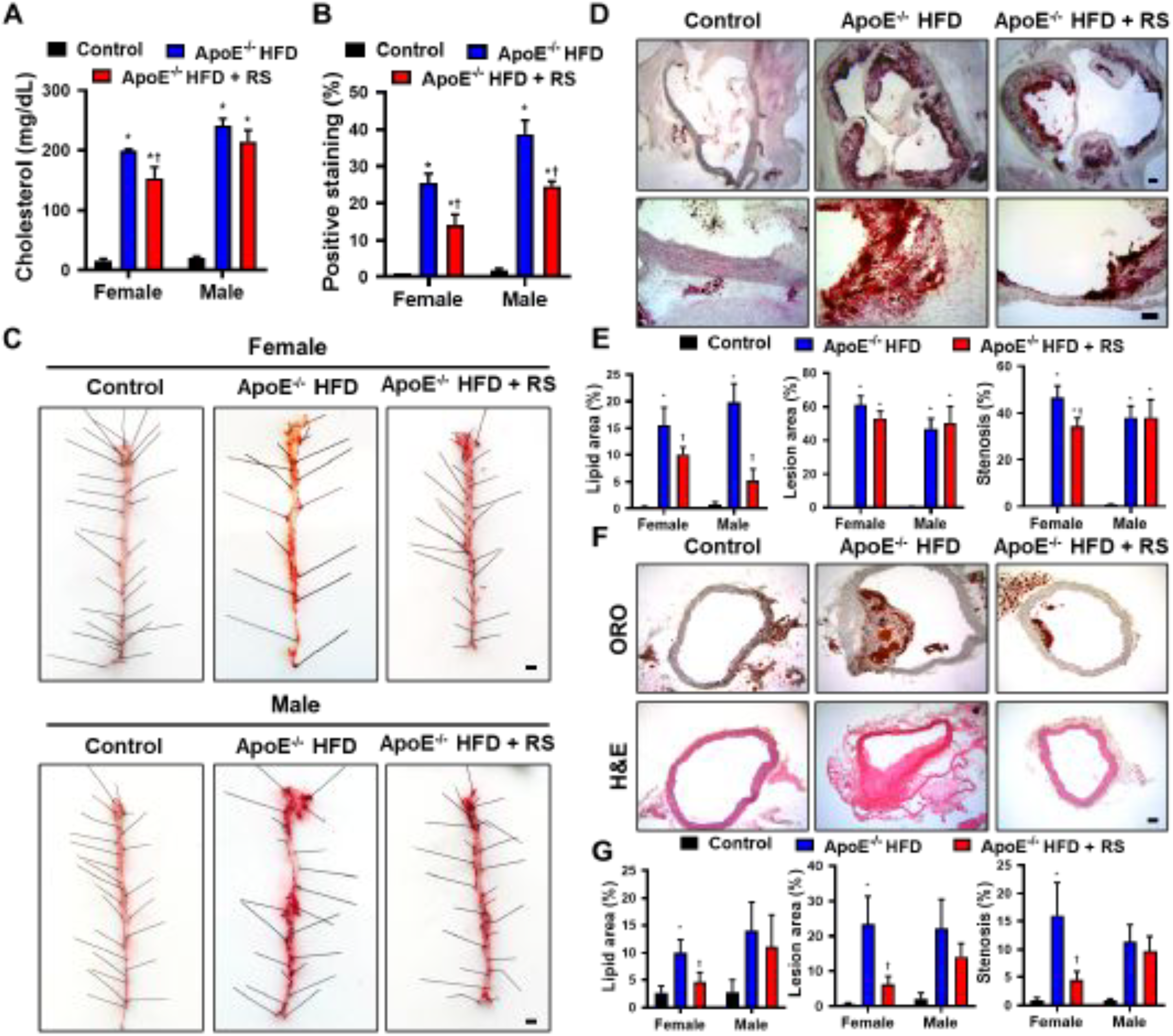
Rhamnan sulfate reduces atherosclerotic plaque area and plasma cholesterol in ApoE^-/-^ mice. (A) Plasma cholesterol was lower by 22.5% in female ApoE^-/-^ mice fed HFD and RS (n = 10). \**p <* 0.05 versus control, ^†^*p <* 0.05 versus ApoE^-/-^ HFD. (B) Lipid deposition in the whole aorta was reduced by 45.2% in female and 36.4% in male ApoE^-/-^ mice fed HFD and RS (n = 3). \**p <* 0.05 versus control, ^†^*p <* 0.05 versus ApoE^-/-^ HFD. (C) En face staining of aortas of female and male mice for C57BL/6 mice fed a standard diet, ApoE^-/-^ mice fed a HFD and ApoE^-/-^ mice fed a HFD with RS. Scale bar = 100 μm. (D) Aortic arch sections of female (shown) and male mice in the three groups were stained with Oil Red-O. Scale bar = 100 μm. (E) Lipid deposition, lesion area and stenosis were reduced in aortic arches of female ApoE^-/-^ mice fed HFD with RS. Lipid deposition only was reduced in aortic arches of male ApoE^-/-^ mice fed HFD with RS (n = 7). \**p <* 0.05 versus control, ^†^*p <* 0.05 versus ApoE^-/-^ HFD. (F) Histological sections of the thoracic aorta of female (shown) and male mice in the three groups were stained with Oil Red-O and hematoxylin and eosin. Scale bar = 100 μm. (G) Lipid deposition, lesion area and stenosis were reduced in thoracic aortas of female ApoE^-/-^ mice fed a HFD with RS (n = 7). \**p <* 0.05 versus control, ^†^*p <* 0.05 versus ApoE^-/-^ HFD.

### Oral rhamnan sulfate does not affect body weight or adipose tissue deposition in ApoE^-/-^ mice

There was no difference in the weight of the mice between the RS treated and untreated mice for both male and female groups (**Supplemental Fig. 6A, B**). There was also no significant difference in the ratio of liver, inguinal white adipose tissue (iWAT) and gonadal white adipose tissue (gWAT) to total body weight in female mice (**Supplemental Fig. 6C**). Also, characterization of the iWAT and gWAT showed no change in size of adipocytes in treated female mice (**Supplemental Fig. 7A**). In male mice, the ratio of gWAT to total body weight increased in mice treated with RS (**Supplemental Fig. 7D**). The size of adipocytes remained the same in both iWAT and gWAT in the HFD and HFD+RS treated groups (**Supplemental Fig. 7B**).

### Rhamnan sulfate treatment reduces high fat diet induced increases in blood velocity in ApoE^-/-^ mice

To monitor plaque development over the course of the experiment, we measured blood velocity in the aorta and left common carotid arteries of ApoE^-/-^ mice. Peak systolic velocity (PSV), end diastolic velocity (EDV) and mean velocity (MV) were calculated in the ascending and descending aorta, aortic arch and, carotid artery. In female mice, PSV, EDV and, MV decreased with RS treatment in the ascending aorta (**Supplemental Fig. 8, Supplemental Fig. 9A**). In the aortic arch for female mice, there was no change in blood flow velocities with RS treatment but there was a significant decrease in flow velocities in the descending aorta for the PSV and EDV with RS treatment (**Supplemental Fig. 8**). All flow velocities decreased in the carotid artery with RS treatment as well for female mice (**Supplemental Fig. 8; Supplemental Fig. 10A**). In male mice, there was no change in all three velocities in the ascending aorta and only EDV decreased in the aortic arch (**Supplemental Fig. 8; Supplemental Fig. 9B**). In the descending aorta, PSV, EDV and, MV all decreased with RS treatment while there no change in the velocities in the carotid artery (**Supplemental Fig. 8; Supplemental Fig. 10B**). In addition, we found that the elasticity, measured by circumferential strain, of the aorta during the cardiac cycle was higher in female mice treated with RS (**Supplemental Fig. 11**).

### Rhamnan sulfate reduces vascular inflammation in female mice but not male mice

To quantify the efficacy of oral RS in reducing vascular inflammation, we examined the presence of macrophages and activation of the NF-κB pathway in the ApoE^-/-^ mice treated with HFD or HFD with RS supplementation. There was a significant decrease in macrophages (F4/80 positive cells) in histological sections from the aortic root in female mice treated with RS in comparison to the HFD group but there was no change in male mice (**Fig. 6A, B**). Also, IκBα, an inhibitor of NF-κB, was significantly upregulated in female mice (**Fig. 6C, D**). Phosphorylation of the p65 subunit of NF-κB was also reduced in female ApoE^-/-^ mice on a HFD treated with RS (**Fig. 6E; Supplemental Fig. 13**). Phosphorylation of p65, IκBα and F4/80 were not affected by RS treatment in male mice (**Fig. 6B, D, E**).

**Figure 6.**
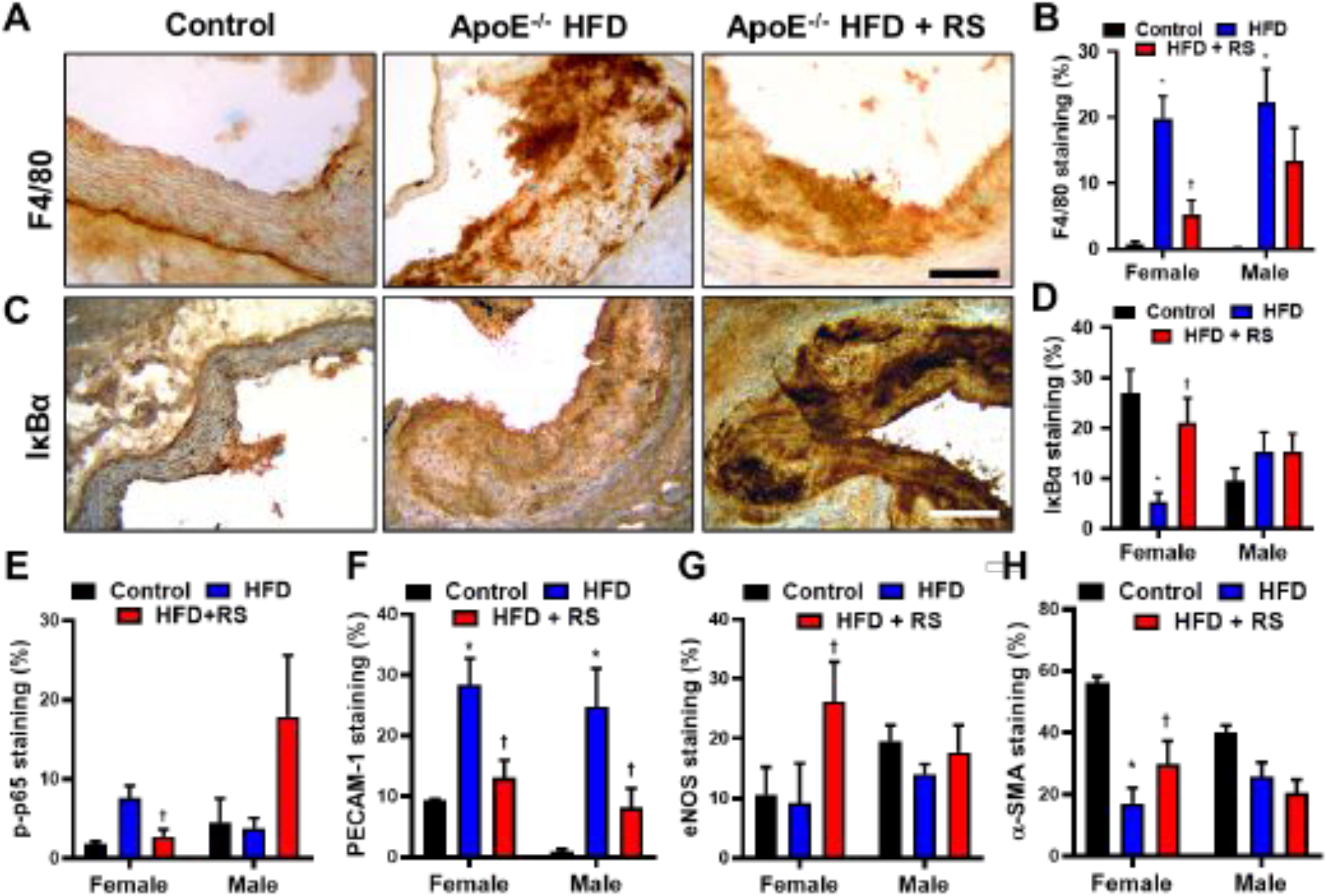
Attenuation of inflammatory markers by RS in the aortic arch. (A) Aortic arch histology sections of female (shown) and male mice in the three groups were stained for F4/80. Scale bar = 100 μm. (B) In female mice treated with RS, there was a reduction of F4/80 positive staining (n = 7). \**p <* 0.05 versus control STD, ^†^*p <* 0.05 versus ApoE^-/-^ HFD. (C) Aortic arch sections were also stained for IκBα. Scale bar = 100 μm. (D) There was an increase in IκBα positive staining in female ApoE^-/-^ mice fed HFD with RS (n = 7). \**p <* 0.05 versus control STD, ^†^*p <* 0.05 versus ApoE^-/-^ HFD. (E) In female mice, phosphorylation of NF-κB/p65 also decreased with RS treatment (n = 7). \**p <* 0.05 versus control STD, ^†^*p <* 0.05 versus ApoE^-/-^ HFD. (F) Positive staining for PECAM-1 was lower in female and male mice treated with RS (n = 7). \**p <* 0.05 versus control STD, ^†^*p <* 0.05 versus ApoE^-/-^ HFD. Finally, in female mice, (G) eNOS and (H) α-SMA staining were increased with RS treatment (n = 7). \**p <* 0.05 versus control STD, ^†^*p <* 0.05 versus ApoE^-/-^ HFD.

### Treatment with RS reduces plaque angiogenesis and enhances endothelial eNOS production in ApoE^*-/-*^ mice on a high fat diet

During atherosclerotic inflammation, endothelial permeability is increased by PECAM-1 upregulation as it promotes leukocyte transmigration and integrin activation. In both female and male ApoE^-/-^ mice, PECAM-1 was decreased with RS treatment (**Fig. 6F; Supplemental Fig. 12**). Impaired eNOS levels lead to atherogenesis though increased nitric oxide breakdown and superoxide production.[42] In female mice treated with RS, there was higher eNOS production in the aortic arch (**Fig. 6G; Supplemental Fig. 12**). There was increased area of α-SMA staining in the aortic arch of female ApoE^-/-^ mice treated with RS, reflecting the reduction in plaque size (**Fig. 6H; Supplemental Fig. 12**). In male mice, eNOS and α-SMA were not affected by RS treatment (**Fig. 6G, H**).

## Discussion

Atherosclerosis is a chronic disease characterized by inflammation, lipid accumulation and the progressive development of plaques.[1],[2] Lipid lowering therapies including statins have provided significant reductions in the risk of fatal coronary heart disease and non-fatal myocardial infarction. However, in spite of these advancements, atherosclerotic disease continues to be a pervasive clinical problem.[43] Atherosclerosis slowly progresses for life, placing more strict requirements on pharmaceutical products to treat vascular disease in comparison to many other diseases. Compounds for treating atherosclerosis will need to be taken orally by patients on a daily basis for decades, requiring long-term safety and representing a major investment by healthcare systems to provide for patients. Our studies demonstrate that rhamnan sulfate has potent anti-inflammatory properties, reduces vSMC proliferation and reduces atherosclerosis in a hyperlipidemic mouse model. As rhamnan sulfate is relatively inexpensive and has been consumed by millions of people as part of their diet, it would present an easily implementable adjuvant therapy to lipid lowering drugs and other treatments for atherosclerotic disease if it were effective in human patients.

Our studies indicate that rhamnan sulfate has atheroprotective effects though multiple modes of action. It reduces migration of vascular cells and binds to pro-growth and inflammatory growth factors with an affinity similar to that of heparin, suppressing the growth of vSMCs. Our studies suggest that this effect could proceed through binding of growth factors including FGF-2 and PDGF-BB. However, RS also inhibits migration in the absence of stimulation with these factors suggesting it may also act by directly altering cellular adhesion/migration mechanisms. The uptake of RS was reduced in both endothelial and vascular smooth muscle cells with rottlerin treatment. This indicates that a major mechanism of entry into the cells is macropinocytosis. In vSMCs, mitomycin treatment also reduced nuclear levels of RS, implying that RS may enter the nucleus during mitosis. This effect may be more pronounced in vSMCs due to their higher proliferation rate.

Our work indicates that rhamnan sulfate accumulates in vascular cells over the course of several days and is also deposited in the extracellular matrix. Interestingly, rhamnan sulfate appears to be able to access and bind to nuclear protein during cell division. This finding suggests a novel mechanism of action for RS, in which it accumulates in the cytoplasm of the cells followed by binding to nuclear proteins, including NF-κB, during cell division. The polyanionic structure of rhamnan sulfate may mimic that of DNA, allowing it to competitively bind to regions of proteins that bind to DNA. In comparison, heparin can facilitate the nuclear entry of growth factors but heparin and its anti-proliferative derivatives do not enter the nucleus of vSMCs.[44] Prior studies have also suggested that the cellular localization of low molecular weight heparin is dependent on sulfation; however, none of the modified forms are found in the nucleus.[45] Thus, rhamnan sulfate is likely not simply acting as a heparin analogue in this activity.

A striking feature of our findings is the differences in response between male and female animals treated with rhamnan sulfate. In the blood vessels in mice treated with rhamnan sulfate, at the aortic root there was a more pronounced reduction in lipid area for male mice, but the overall lesion and stenotic response was only significantly lower in females. In the thoracic aorta, only female mice had a reduction in plaque size and stenosis. We also observed a stronger cholesterol-lowering effect of rhamnan sulfate in female mice compared to male mice. Lipid lowering effects have been observed for marine polysaccharides including fucoidan, ulvan, and laminarin sulfate.[46-48] In comparison to these, rhamnan sulfate did not produce as dramatic reductions in lipids. The anti-inflammatory properties of rhamnan sulfate appeared to be stronger in female mice as well, including significant lowering of plaque inflammation and lipids with rhamnan sulfate treatment.

Overall, our studies demonstrate that rhamnan sulfate can provide significant benefits for inhibiting the development of atherosclerosis and acts though multiple mechanisms including the reduction of inflammation, inhibition of vascular cell proliferation and enhanced endothelial barrier function. Due to the low cost and ease of availability, RS may have high potential as an easily implementable adjunct therapy for atherosclerosis. Our studies used a highly purified form of rhamnan sulfate, which may allow it to have increased binding capacity and activity in comparison to less pure forms. Further studies would be needed to determine if less purified forms of the compound have similar activity and if particular subfractions of RS can be linked to specific mechanistic activities.

## Acknowledgements

The authors gratefully acknowledge funding though the American Heart Association (17IRG33410888), the DOD CDMRP (W81XWH-16-1-0580; W81XWH-16-1-0582) and the National Institutes of Health (1R21EB023551-01; 1R21EB024147-01A1; 1R01HL141761-01) to ABB. This work was funded by National Institutes of Health Grants DK111958, CA231074, NS088496 and AG062344 to RJL.

## Disclosures

None.

---

**Supplemental Table 1.** Primary Antibodies Used for Immunoblotting.

| **Target Protein** | **Company** | **Catalog #** | **Species/Isotype** | **Dilution Ratio** |
| --- | --- | --- | --- | --- |
| GAPDH | Cell Signaling | 2118S | anti-rabbit | 1:500 |
| α-SMA | Abcam | ab21027 | anti-rabbit | 1:500 |
| α-SMA (Cy3) | Sigma-Aldrich | C6198 | Anti-mouse | 1:200 |
| α-Tubulin | Sigma-Aldrich | T9026 | anti-mouse | 1:500 |
| β-Actin | Santa Cruz Biotech | sc-47778 | anti-mouse | 1:500 |
| eNOS | Santa Cruz Biotech | sc-654 | Anti-rabbit | 1:500 |
| F4/80 | Abcam | Ab6640 | anti-mouse | 1:500 |
| FAK | Cell Signaling | 3285S | Anti-rabbit | 1:100 |
| Histone H3 | Cell Signaling | 9715S | anti-mouse | 1:500 |
| IκBα | Cell Signaling | 4814S | anti-rabbit | 1:500 |
| IKKα | Santa Cruz Biotech | sc-7606 | anti-mouse | 1:500 |
| IKKβ | Cell Signaling | 2678S | anti-rabbit | 1:500 |
| NF-κBp65 | Thermo Fisher Scientific | PA1186 | anti-rabbit | 1:500 |
| PECAM-1 | Cell Signaling | 3528S | anti-mouse | 1:500 |
| Phospho-IκBα (S32/34) | Thermo Fisher Scientific | MA515224 | anti-rabbit | 1:500 |
| Phospho-IKKα/$\beta$ | Cell Signaling | 2797S | Anti-rabbit | 1:500 |
| TKS5 | Sigma-Aldrich | 09-403 | Anti-rabbit | 1:100 |

**Supplemental Table 2.** Rhamnan Sulfate Binding Affinity.

| **Interaction** | **k<sub>a</sub> (ms<sup>-1</sup>)** | **k<sub>d</sub> (s<sup>-1</sup>)** | **K<sub>D</sub> (M)** |
| --- | --- | --- | --- |
| PDGF-BB to RS | 2.5 x 10<sup>5</sup><br>(±4.9 x 10<sup>3</sup>) | 5.4 x 10<sup>-3</sup><br>(±1.1 x 10<sup>-3</sup>) | 2.8 x 10<sup>-8</sup> |
| PDGF-BB to Heparin | 2.6 x 10<sup>5</sup><br>(±4.3 x 10<sup>3</sup>) | 4.9 x 10<sup>-3</sup><br>(±8.3 x 10<sup>-5</sup>) | 1.9 x 10<sup>-8</sup> |
| FGF-2 to RS | 3.0 x 10<sup>5</sup><br>(±6.0 x 10<sup>3</sup>) | 7.7 x 10<sup>-3</sup><br>(±8.1 x 10<sup>-5</sup>) | 2.6 x 10<sup>-8</sup> |
| FGF-2 to Heparin | 5.9 x 10<sup>5</sup><br>(±1.4 x 10<sup>4</sup>) | 5.3 x 10<sup>-3</sup><br>(±8.1 x 10<sup>-5</sup>) | 9.0x 10 |
| NFκB p50 to RS | 1.8 x 10<sup>4</sup><br>(±721) | 1.9 x 10<sup>-3</sup><br>(±7.9 x 10<sup>-5</sup>) | 1.1 x 10<sup>-7</sup> |
| NFκB p50 to Heparin | 1.4 x 10<sup>4</sup><br>(±627) | 2.1 x 10<sup>-3</sup><br>(±8.4 x 10<sup>-5</sup>) | 1.5 x 10<sup>-7</sup> |
| NFκB p65 to RS | 4.8 x 10<sup>5</sup><br>(±1.1 x 10<sup>4</sup>) | 0.011<br>(±2.2 x 10<sup>-4</sup>) | 2.3 x 10<sup>-8</sup> |
| NFκB p65 to Heparin | 2.6 x 10<sup>5</sup><br>(±5.2 x 10<sup>5</sup>) | 0.011<br>(±3.4 x 10<sup>-4</sup>) | 4.2 x 10<sup>-8</sup> |

## Supplemental Figure Legends

**Supplemental Figure 1.**
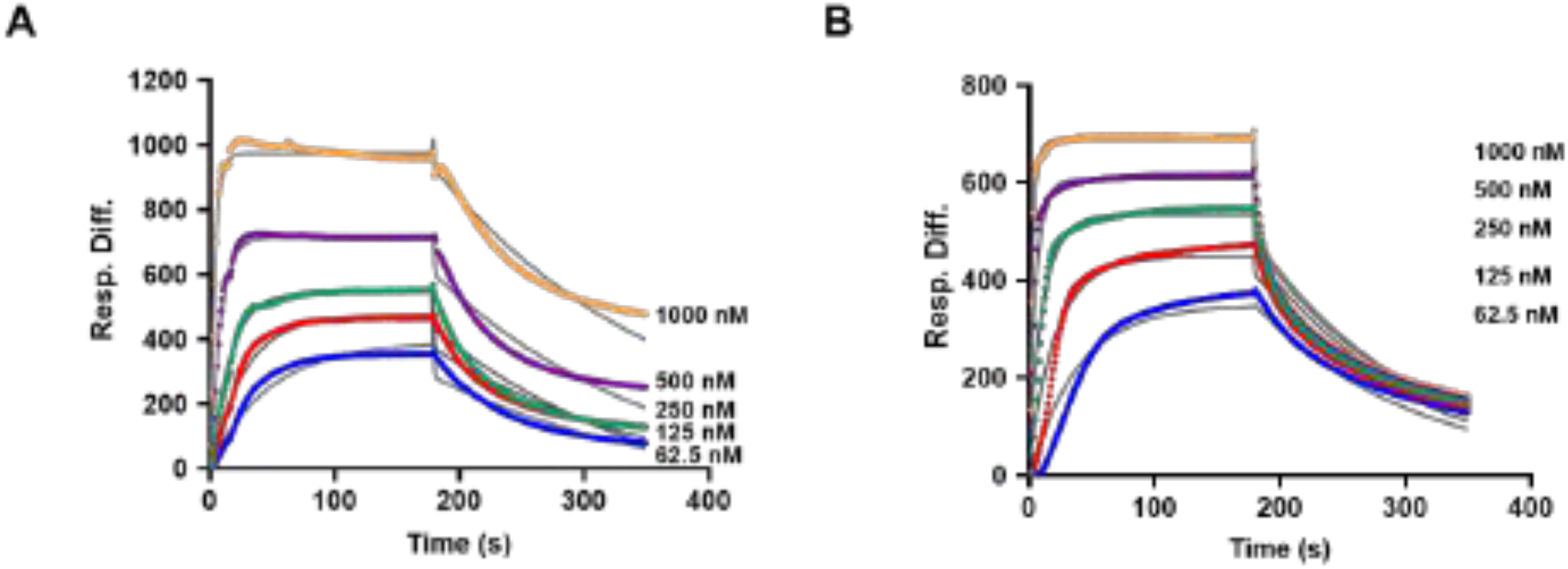
Dilution of (A) PDGF-BB and (B) FGF-2 were introduced on a sensor surface with RS and dissociation with HBS-EP buffer measured for three minutes. After three minutes, the surface was regenerated with 2 M NaCl and total response monitored in sensorgram (n = 5).

**Supplemental Figure 2.**
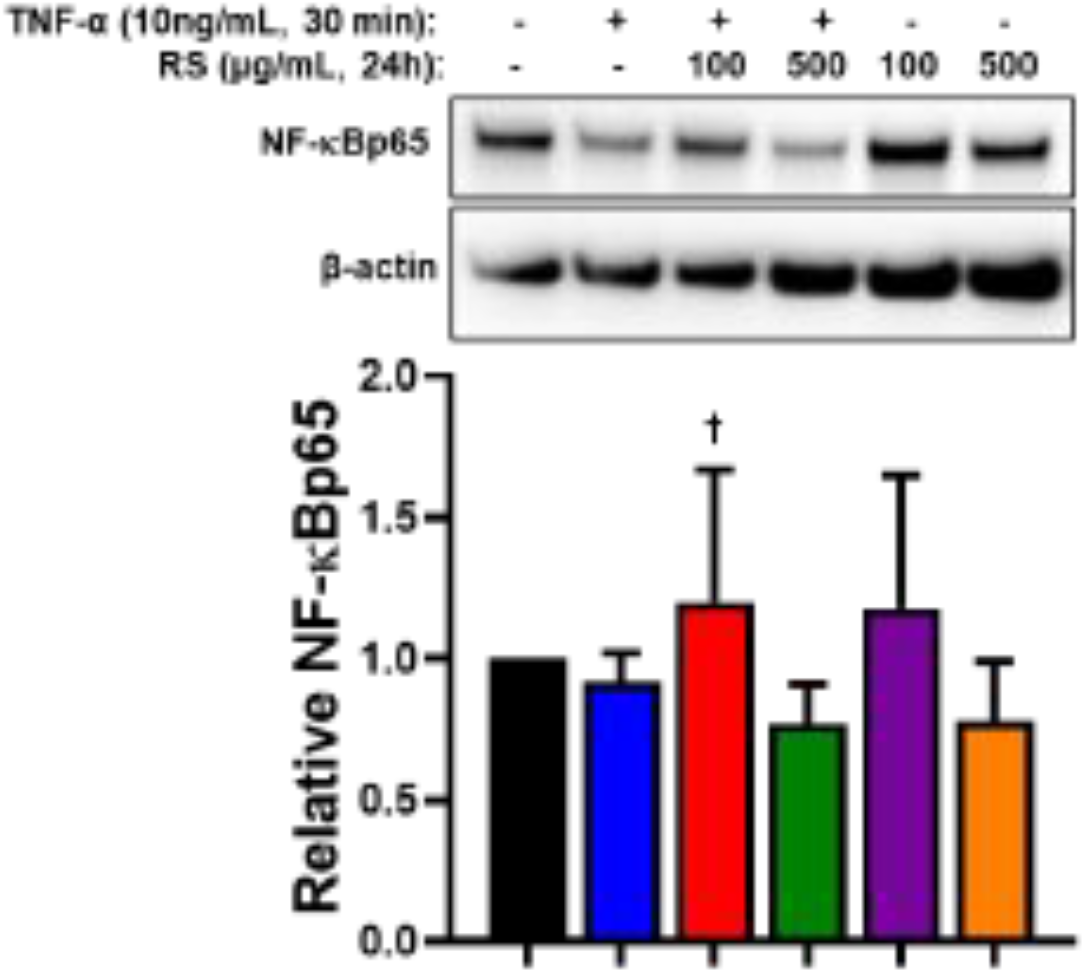
Western blot analysis of cytosol fraction of p65 subunit of NF-κB. Cytosolic protein increased with 100 µg/mL RS dose but there was no change with other treatments (n = 8). ^†^*p <* 0.05 versus TNF-α.

**Supplemental Figure 3.**
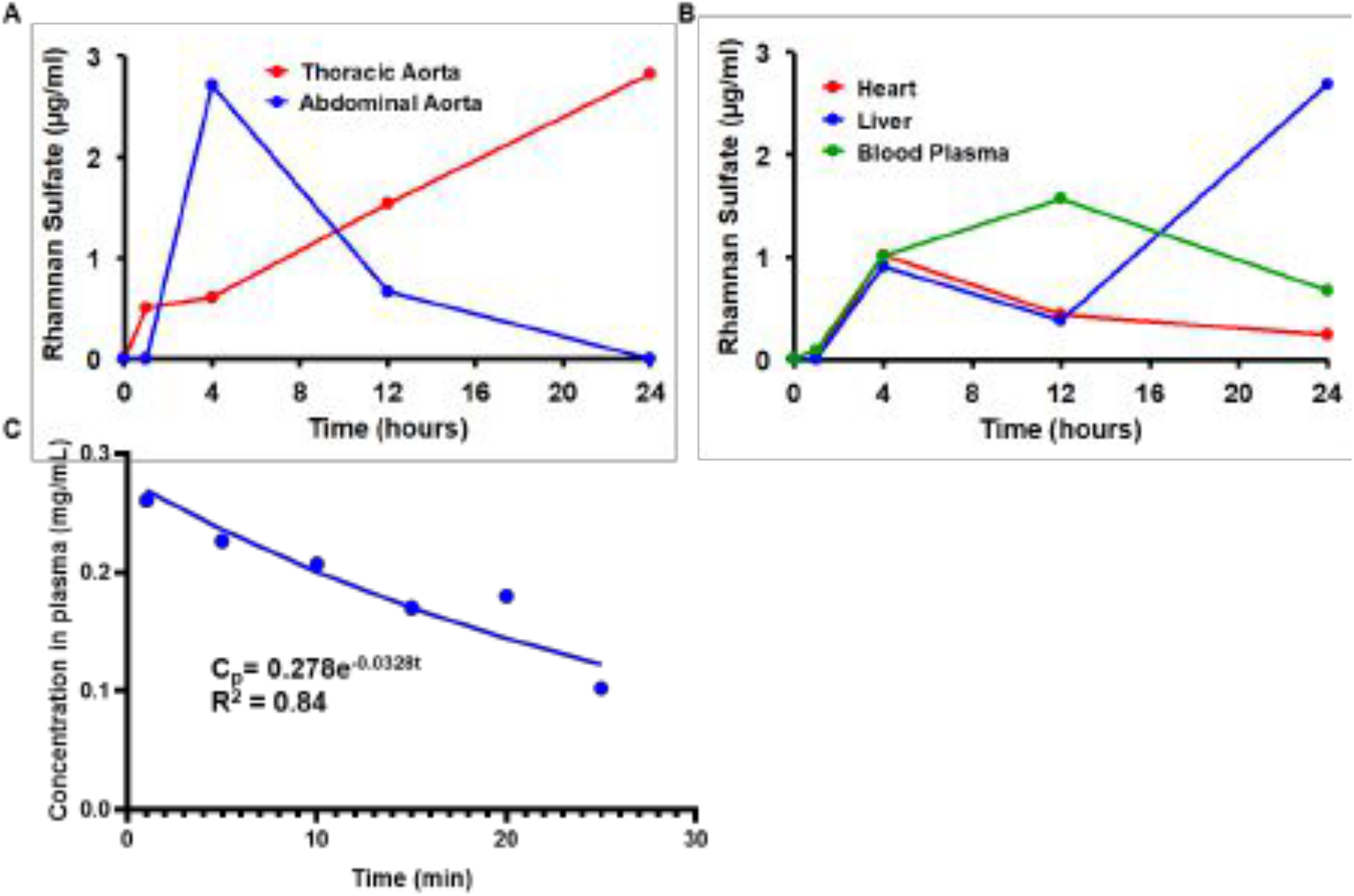
Pharmacokinetics of RS *in vivo*. (A) Rhamnan sulfate concentration decreased after 4 hours in the abdominal aorta but accumulated in the thoracic aorta up to 24 hours after oral gavage (n = 2). (B) Concentration of RS in total blood plasma increased up to 12 hours and then started decreasing. In the liver, there was an initial influx of RS up to 4 hours after oral gavage and then steady accumulation after 12 hours. In the heart, RS concentration peaked at 4 hours after oral gavage (n = 2). (C) Following an intravenous injection, the decay constant of RS was calculated to be 0.0328 min^-1^ and a half-life of 21.13 minutes (n = 3).

**Supplemental Figure 4.**
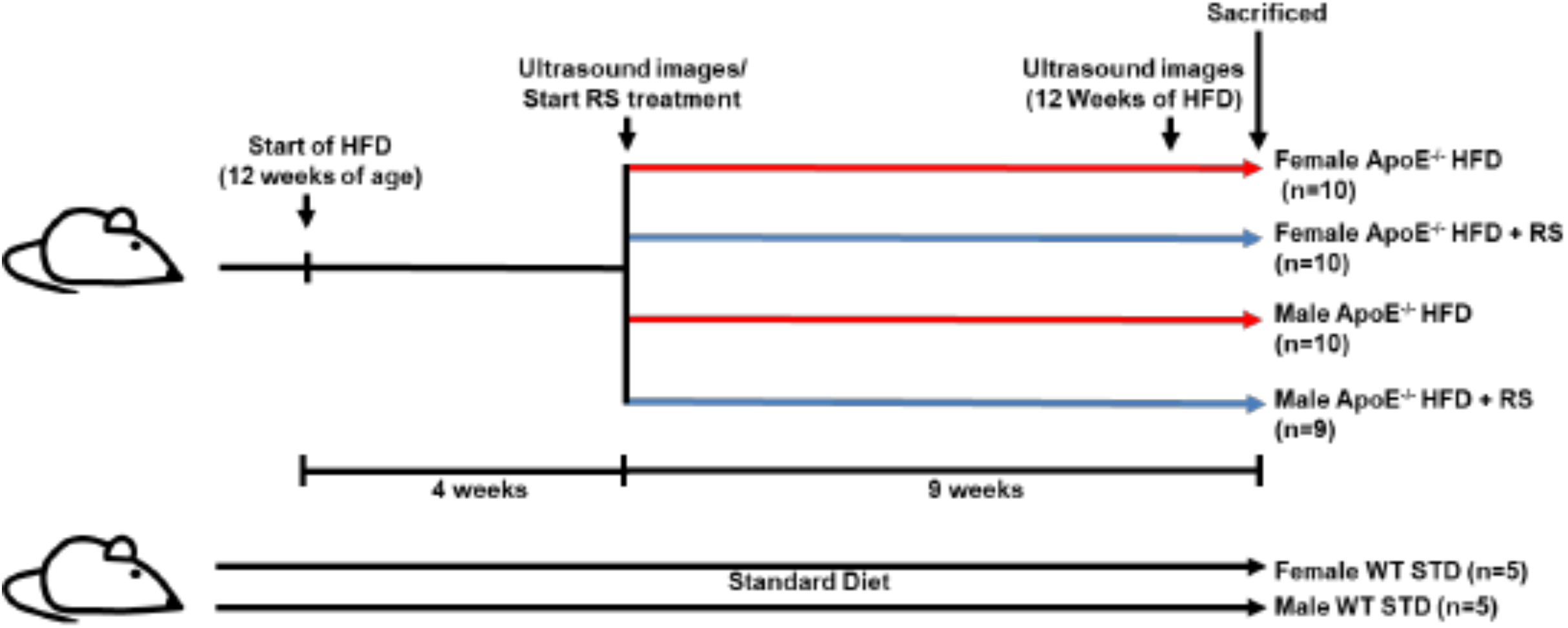
Schematic representation of *in vivo* experiment for prevention of atherosclerosis.

**Supplemental Figure 5.**
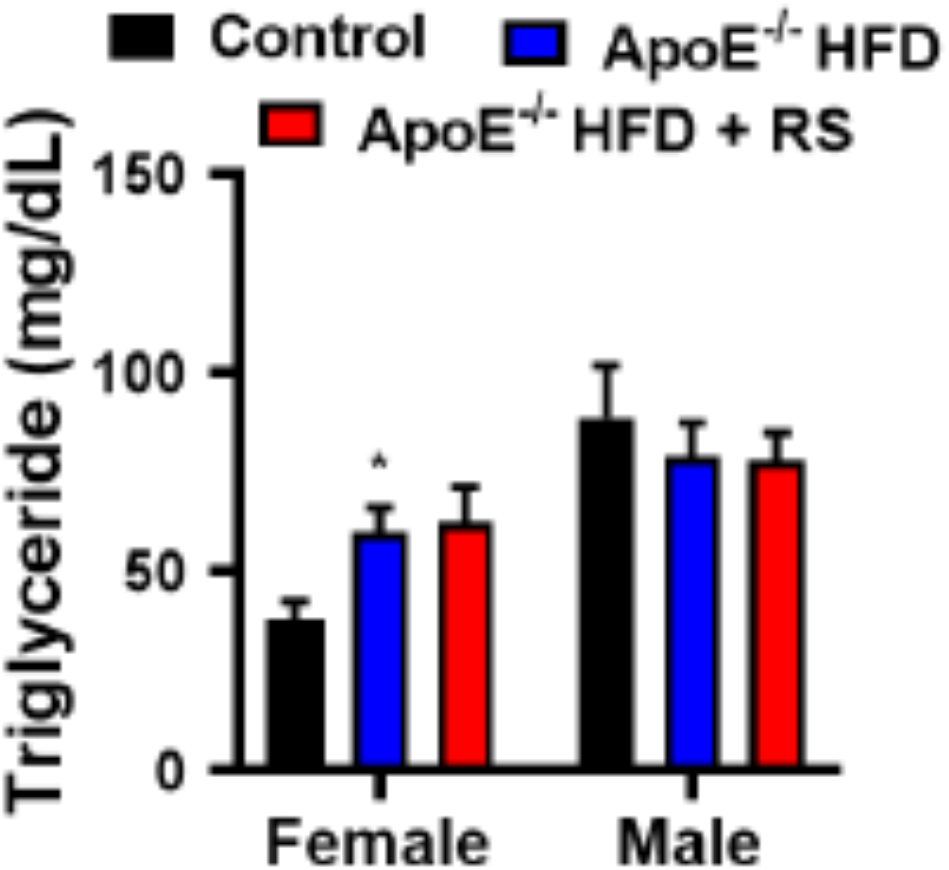
Plasma triglyceride concentration of ApoE^-/-^ mice on HFD. Triglycerides in plasma of female mice increased with HFD but no change with RS treatment for both male and female mice (n = 10).

**Supplemental Figure 6.**
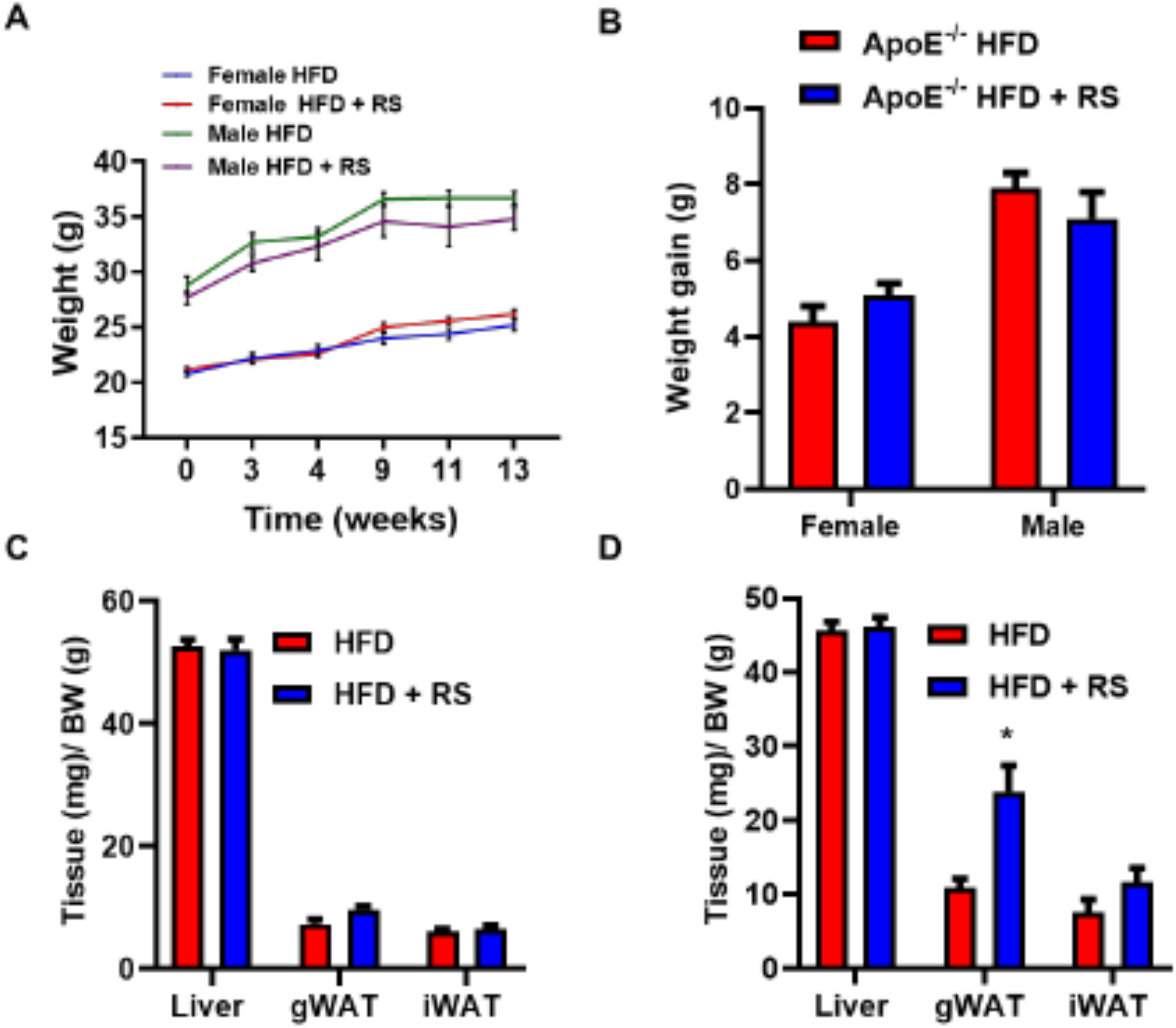
Body and tissue weight of ApoE^-/-^ mice on HFD and RS treatment. (A) Average weights of male and female mice increased over 13 weeks of HFD but there was no significant difference between HFD and HFD+RS groups (n = 10). (B) After 13 weeks, there was no significant change in weight in male and female mice with RS treatment (n = 10). (C) There was no difference in the ratio of average liver, gWAT or iWAT weight to total body weight in female mice (n = 10). (D) In male mice, gWAT to body weight ratio increased with RS treatment but there was no change in the liver or iWAT ratios (n = 10). \**p <* 0.05 versus HFD.

**Supplemental Figure 7.**
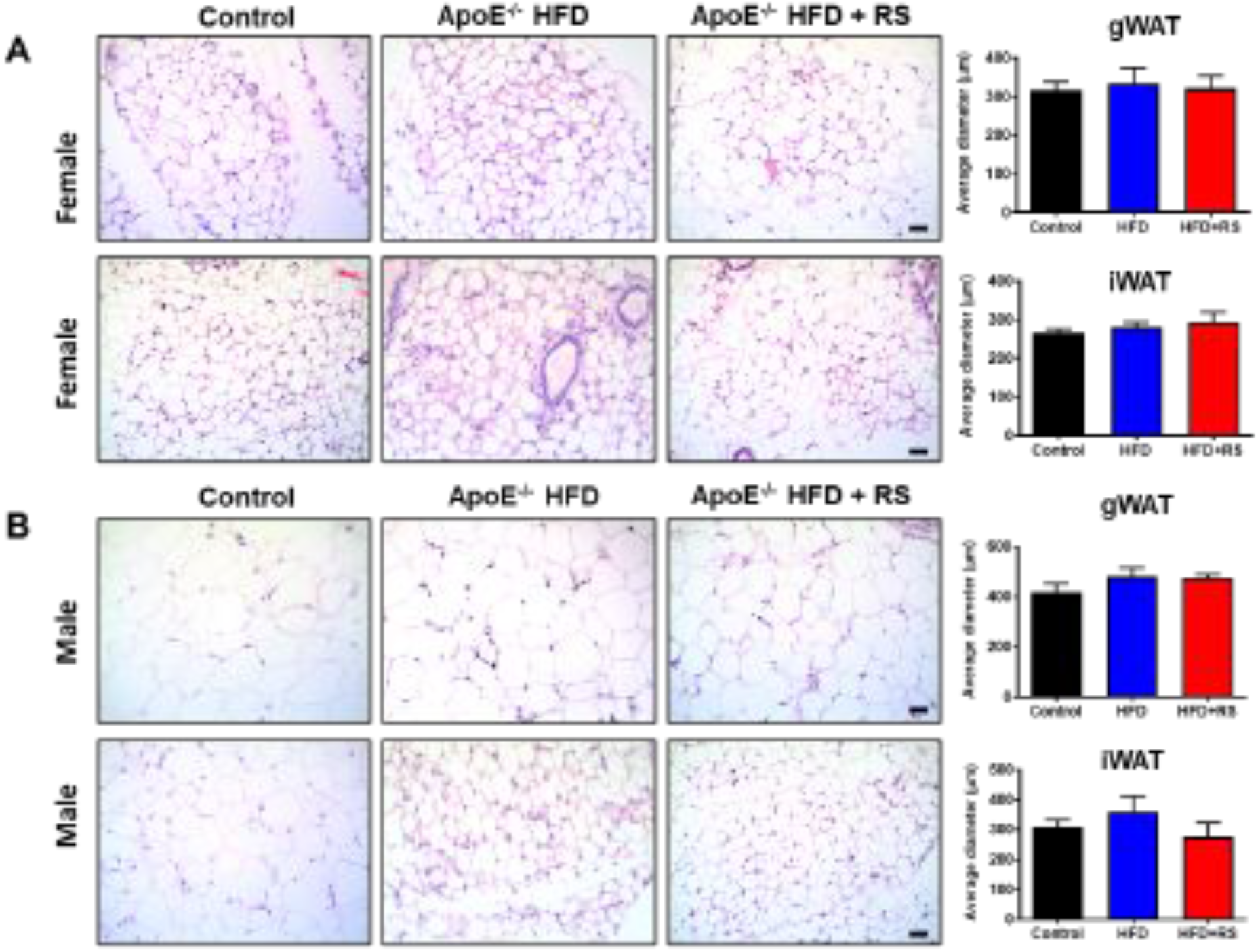
Adipocytes in white adipose tissue are not affected by RS treatment. Histology sections of (A) gWAT and (B) iWAT from female mice were stained with hematoxylin and eosin. There was no change in size of adipocytes with HFD or RS treatment (n = 10). Scale bar = 200 μm. Sections of (C) gWAT and (D) iWAT from male mice were also stained with hematoxylin and eosin. There was no change in size of adipocytes with HFD or RS treatment (n = 10). Scale bar = 200 μm.

**Supplemental Figure 8.**
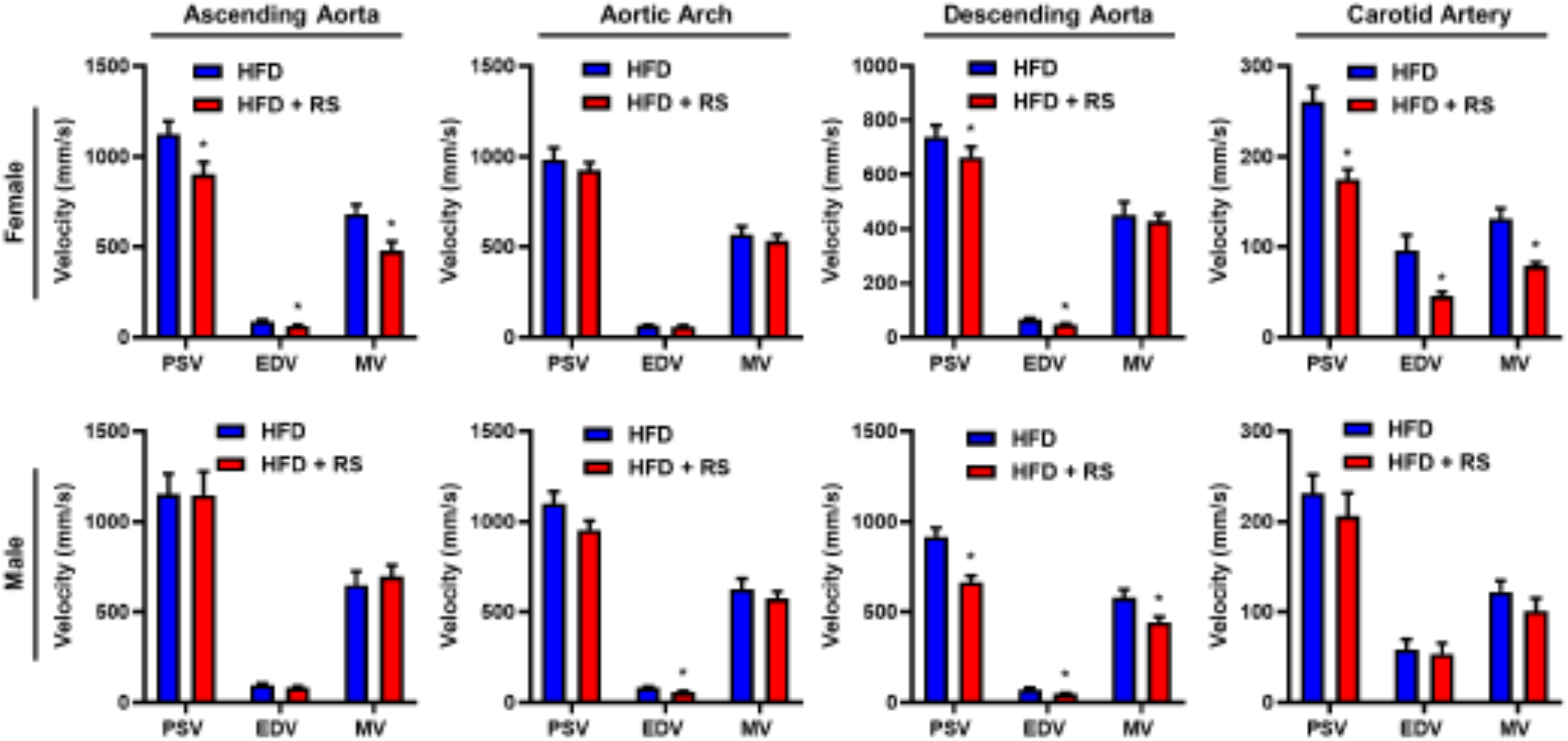
Blood velocity measured by ultrasound is decreased by RS in ApoE^-/-^ mice. After 12 weeks of HFD and RS, PSV, EDV and MV are lower compared to no treatment in ascending aortas of female mice while there was no change in blood velocity in aortic arches. Compared to no treatment, PSV and EDV were lower in descending aortas of female mice. In the carotid arteries, PSV, EDV and MV were lower compared to no treatment. In male mice, there was no change in blood velocity in the ascending aortas while only EDV was lower compared to the HFD group in the aortic arches. In the descending aortas of male mice, PSV, EDV and MV were lower compared to no treatment. There was no change in blood velocity in the carotid arteries of male mice (n = 10). \**p <* 0.05 versus HFD.

**Supplemental Figure 9.**
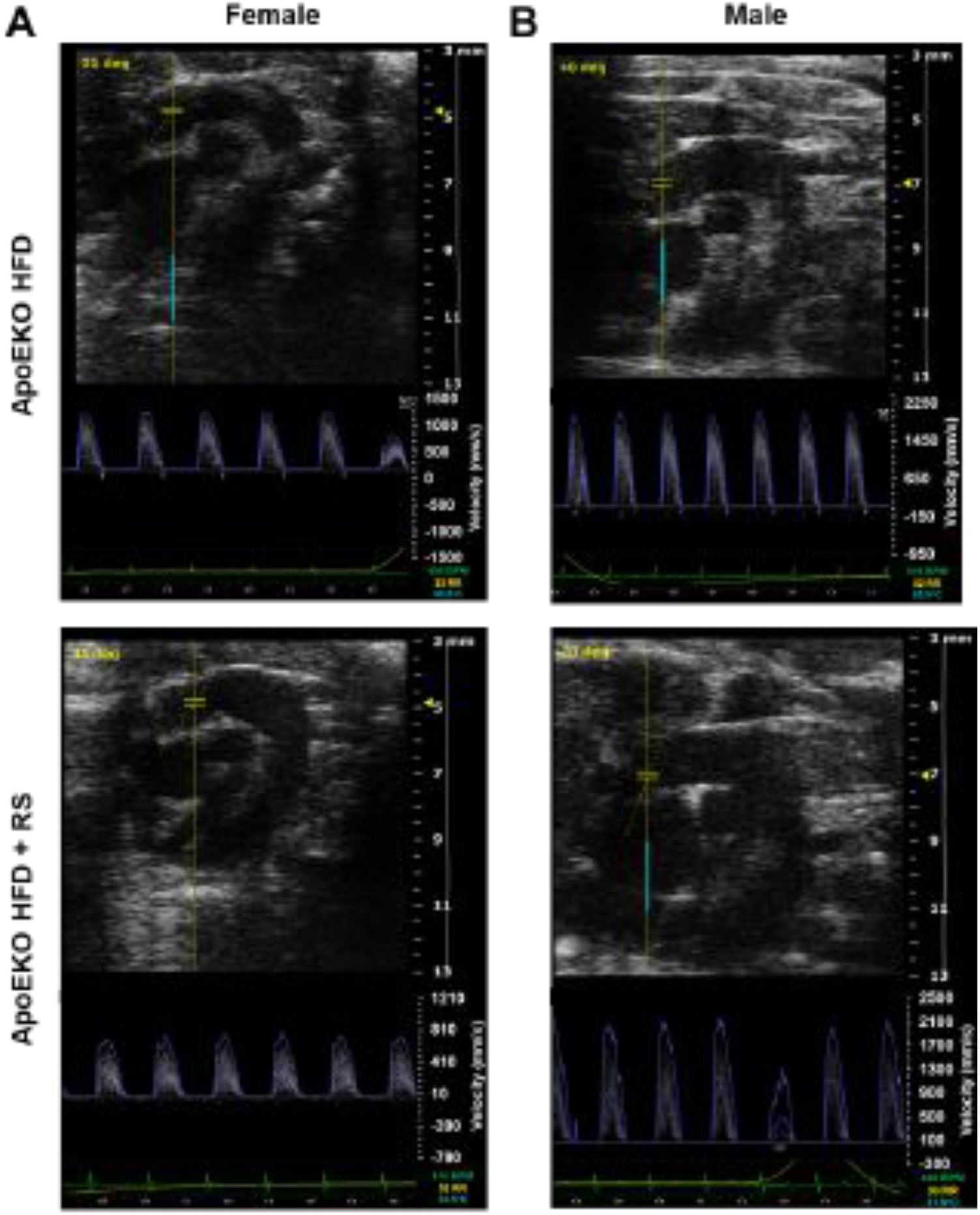
Doppler ultrasound imaging to measure velocity of blood in aorta of ApoE^-/-^ mice. (A) Images and velocity profiles of female ApoE^-/-^ mice fed HFD and HFD+RS (n = 10). (B) Images and velocity profiles of male ApoE^-/-^ mice fed HFD and HFD+RS (n = 10).

**Supplemental Figure 10.**
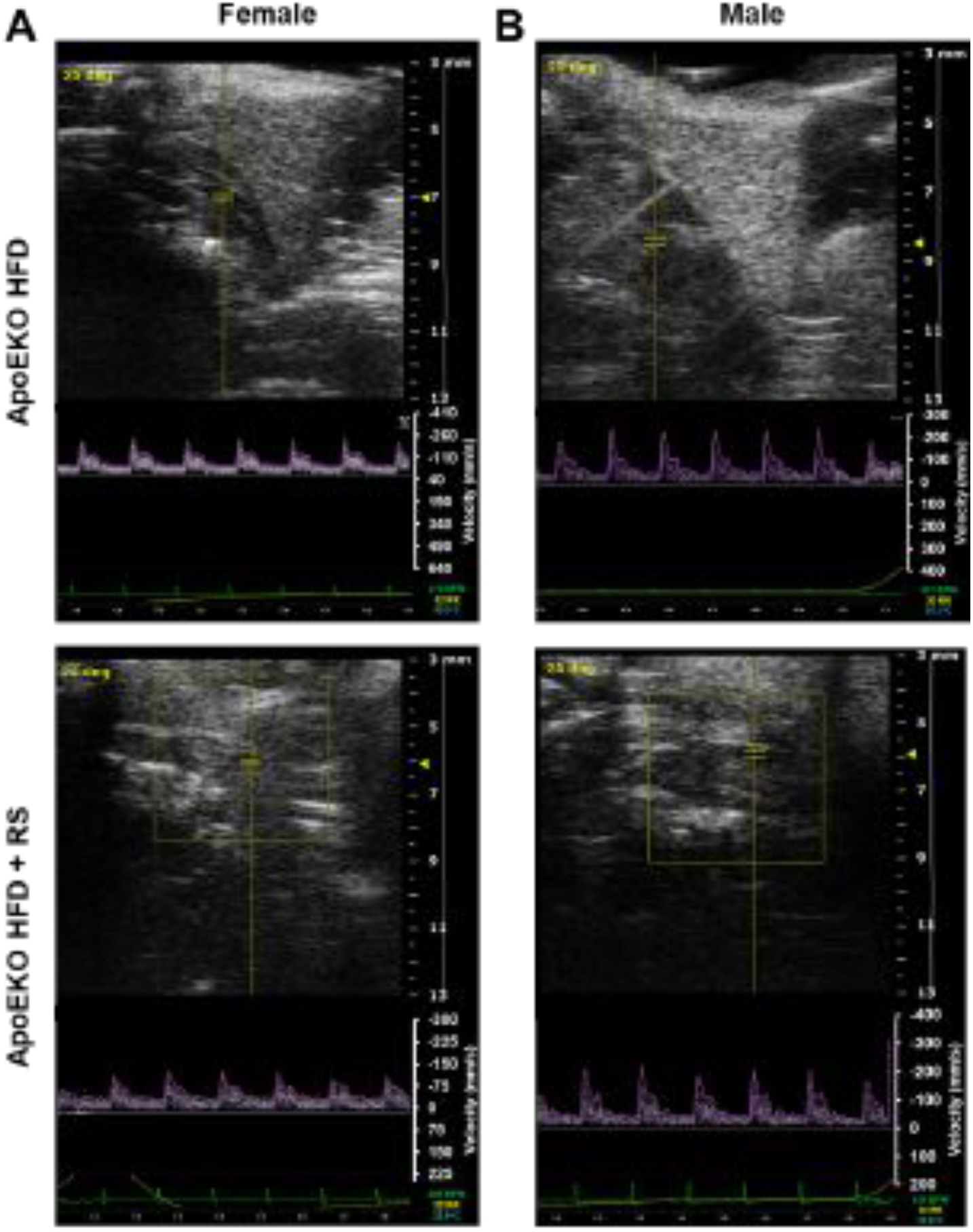
Doppler ultrasound imaging to measure velocity of blood in left common carotid artery of ApoE^-/-^ mice. (A) Images and velocity profiles of female ApoE^-/-^ mice fed HFD and HFD+RS (n = 10). (B) Images and velocity profiles of male ApoE^-/-^ mice fed HFD and HFD+RS (n = 10).

**Supplemental Figure 11.**
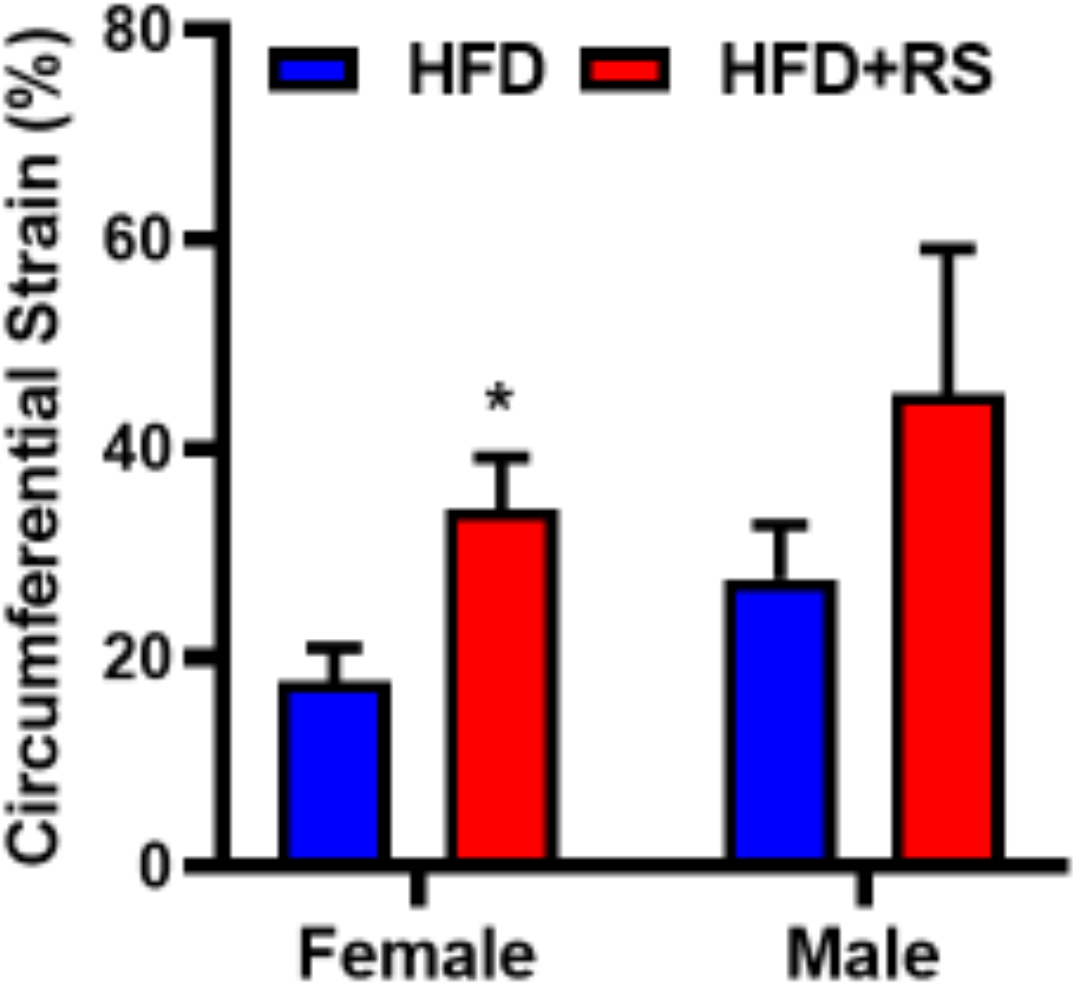
Elasticity of aorta. Elasticity measured by circumferential strain is higher in female ApoE^-/-^ mice treated with RS (n = 10).

**Supplemental Figure 12.**
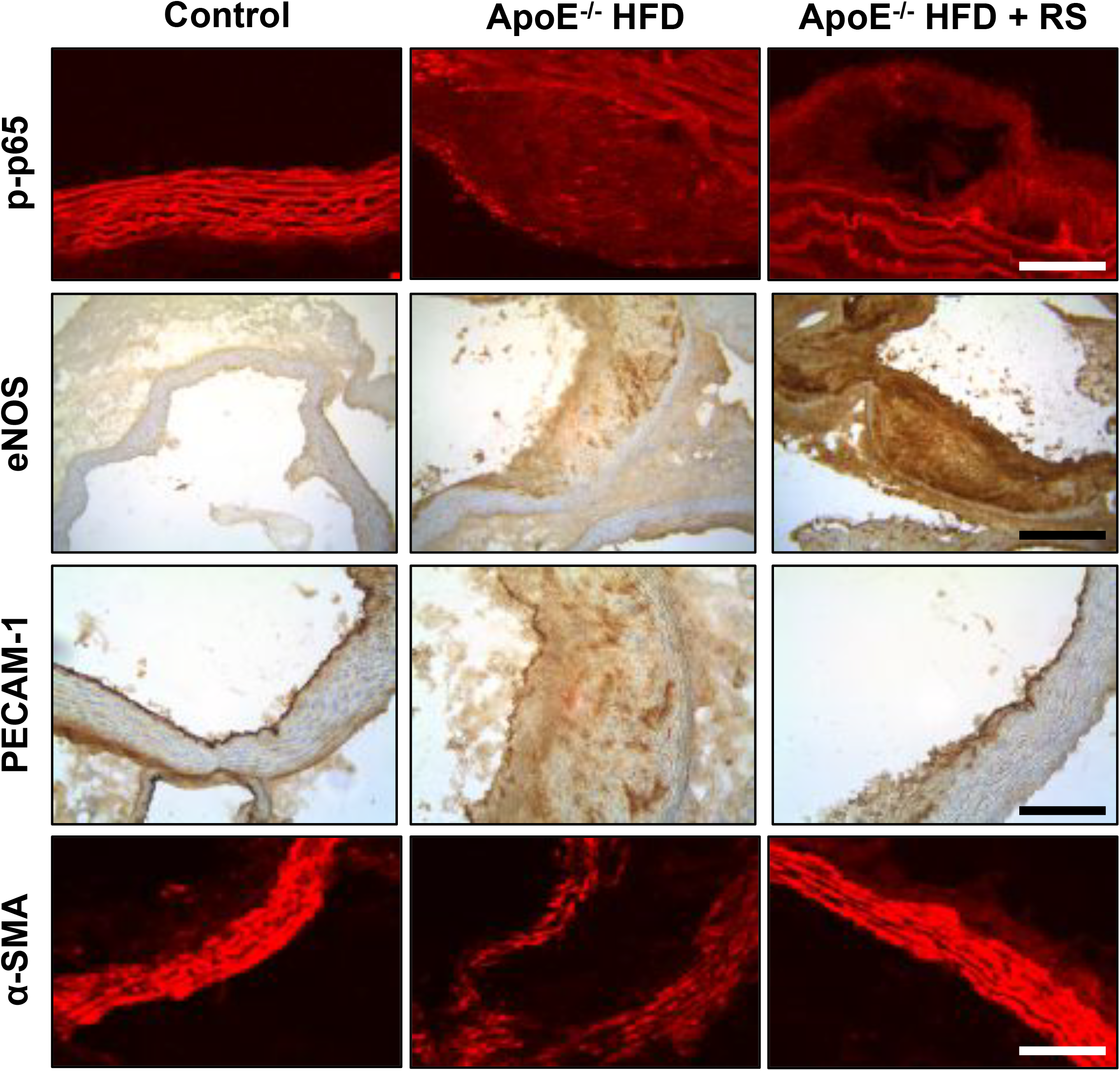
Immunohistochemical staining images of aortic arch from female mice. (A) Immunofluorescent images of NF-κB/p65 staining from the three groups. Scale bar = 20 μm. Images of (B) PECAM-1 and (C) eNOS staining from the three groups. Scale bar = 100 μm. (D) Immunofluorescent images of α-SMA staining from the three groups. Scale bar = 20 μm.

**Supplemental Figure 13.**
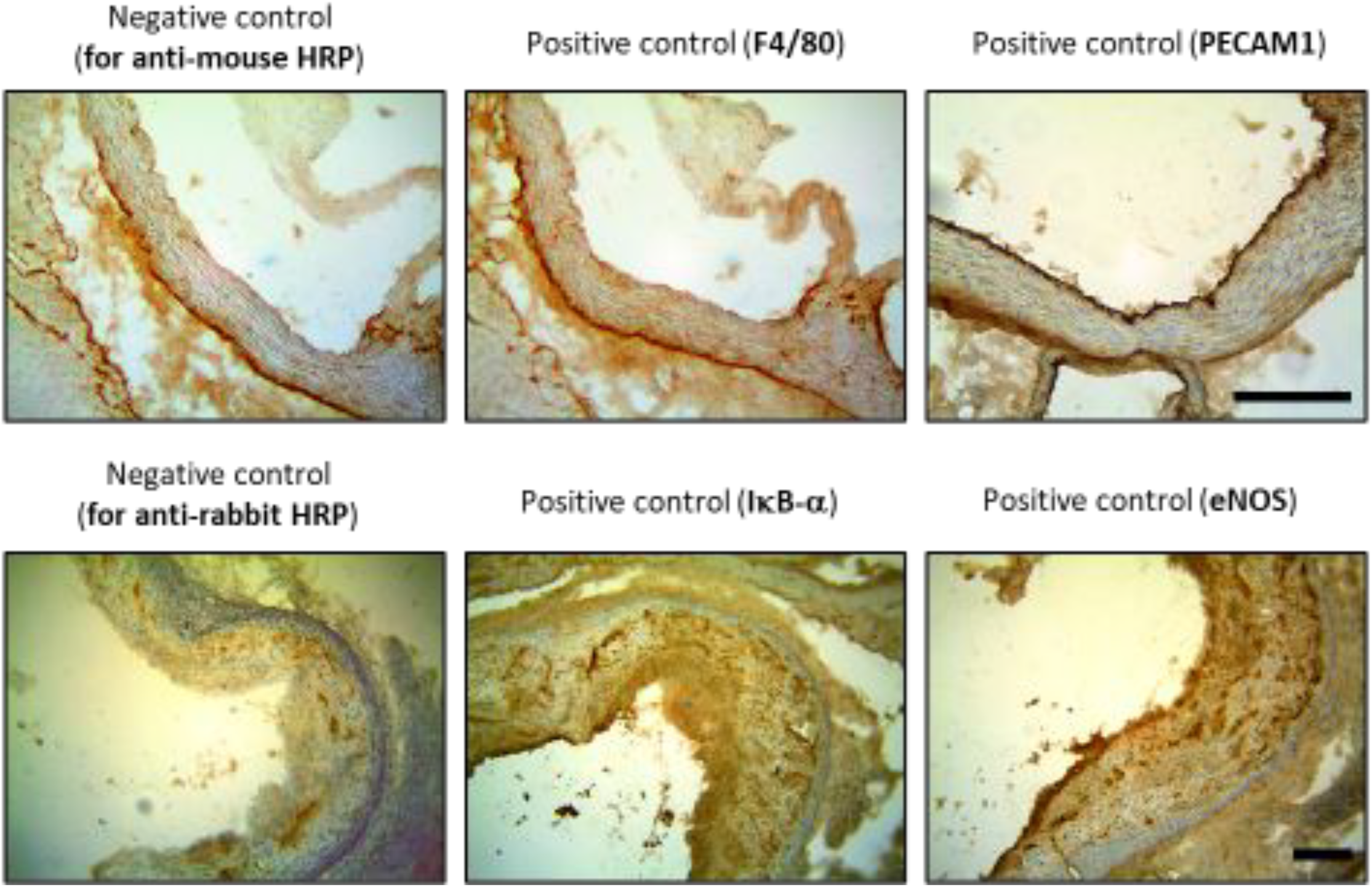
Negative and positive controls of immunohistochemistry staining. Scale bar = 100 μm.

**Supplemental Figure 14.** Volcano plots for comparisons between the groups.

**Supplemental Figure 15.** Clustering analysis for the comparisons between the groups.

**Supplemental Figure 16.** Gene ontology analysis for (A) male or (B) female mice comparing HFD to HFD + RS groups.

## Notes

### Competing Interest Statement

The authors have declared no competing interest.

